# Hippocampal functional connectivity changes associated with active and lecture-based physics learning

**DOI:** 10.1101/2025.09.22.677908

**Authors:** Donisha D. Smith, Jessica E. Bartley, Julio A. Peraza, Katherine L. Bottenhorn, Michael C. Riedel, Taylor Salo, Robert W. Laird, Shannon M. Pruden, Matthew T. Sutherland, Eric Brewe, Angela R. Laird

**Author notes:** Corresponding Author*: Donisha D. Smith, Department of Psychology, Florida International University, Miami, FL, USA.

## Abstract

Introductory university physics courses often face the dual challenge of introducing students to new physics concepts while also addressing their preconceived notions that conflict with scientific principles. Active learning pedagogical approaches, which employ constructivist principles and emphasize active participation in the learning process, have been shown to be effective in teaching complex physics concepts. However, while the behavioral effects of constructivist methodologies are largely understood, the neurobiological underpinnings that facilitate this process remain unclear. Using functional magnetic resonance imaging (fMRI), we assessed students enrolled in either an active learning or lecture-based physics course before and after a 15-week semester of learning and examined changes in hippocampal whole-brain connectivity. We focused on the hippocampus given its critical role in learning and memory. Our findings revealed that hippocampal connectivity with brain regions in the frontal and parietal lobes decreased over time, regardless of instructional approach. Results also indicated that active learning students exhibited increased hippocampal connectivity with parietal, cerebellar, and frontal regions, reflecting experiential learning based on constructivist principles, whereas lecture-based students exhibited increased hippocampal connectivity with the fusiform gyrus, suggesting learning through passive observation. Our findings demonstrate that while some aspects of hippocampal functional connectivity may decrease over time, active vs. passive learning may preferentially enhance hippocampal connectivity during physics learning.

## Introduction

University-level introductory physics is a foundational scientific course that teaches the essential knowledge required to explore more advanced topics in physics, engineering, and related disciplines. In introductory physics, students often encounter formal presentations of physics concepts for the first time <u>(Shafer et al., 2024)</u> and their initial experiences in these courses significantly influence how they engage with future physics-related content <u>(Porter et al., 2024)</u>. Unlike many introductory courses, students in physics courses often arrive with schemas and preconceived mental representations of physics concepts formed through everyday experiences <u>(Chu et al., 2008)</u>. However, research findings suggest that these naive mental schemas are often contradictory to core physics principles. Consequently, introductory physics courses often face the challenge of both delivering content effectively while simultaneously guiding students toward more accurate mental representations that align with core physics principles. To address these challenges, research into physics education has examined the efficacy of various instructional methodologies designed not only to convey complex and counterintuitive concepts, but also to enhance overall understanding of these phenomena. Findings suggest that active learning approaches, which emphasize student-led direct engagement with learning materials and collaborative problem solving, have demonstrated success in addressing these challenges compared to lecture-based approaches, which employ traditional, instructor-led learning methods that involve minimal contributions from students during the instructional process <u>(Dancy et al., 2024;</u> <u>Sandrone et al., 2021)</u>.

From a behavioral perspective, the long-term benefits of active learning are both multifaceted and well understood. Studies assessing active learning approaches to teaching science, technology, mathematics, and engineering (STEM) courses show that this pedagogy is significantly associated with higher examination scores and improved course retention rates <u>(Freeman et al., 2014)</u>. In the context of physics education, active learning typically fosters a deeper and more integrated understanding of physics principles and increases the likelihood that students will develop scientifically informed mental representations of physics concepts, which ultimately results in the development of adaptable problem solving approaches towards general physics-related problems <u>(Hefter et al., 2022; Kotsis, 2024; Roche et al., 2017; Sokoloff et al., 2007)</u>. Conversely, despite our robust understanding of the behavioral benefits of active learning, the neurobiological mechanisms underlying these improvements remain poorly understood. Understanding the neurobiological underpinnings driving the behavioral differences between active and lecture-based learning may aid educational practitioners in refining and developing more effective instructional approaches to improve the efficacy of introductory physics courses at the university level.

The efficacy of active learning approaches may lie in its theoretical roots of constructivist learning theory. Constructivism posits that knowledge acquisition is an active process, where, through diverse experiences, learners can reflect on their understanding and form new connections between new information and prior knowledge, resulting in more robust and integrative schemata of abstract concepts <u>(Pande & Bharathi, 2020)</u>. This suggests a bidirectional relationship wherein experiences contribute to the development of better foundational understanding, while improved foundational understandings allows for more sophisticated associations to be drawn from new experiences. This demonstrates how improved prior knowledge ultimately culminates in more enhanced learning. Furthermore, active learning approaches often apply constructivism through experiential learning <u>(Mughal & Zafar, 2011)</u>, especially when employed in physics courses, where students engage in hands-on experimentation and collaborative problem solving <u>(Brewe et al., 2018; Johari & Muslim, 2018; Md Rashid et al., 2024; Robinson et al., 2016)</u>. This process integrates information from multiple sensory modalities during the learning process (e.g., visual, auditory, sensorimotor) and encourages students to make connections between information obtained from these concrete experiences and abstract physics concepts. The neurobiological processes underlying constructivist learning principles may be facilitated by the hippocampus, a structure known for its central role in integrating new information with existing knowledge structure <u>(Bein et al., 2020)</u> and relational memory which involves associating different elements of information into the same episodic memory <u>(Borders et al., 2017)</u>.

The hippocampus, located in the inner temporal lobe <u>(Anand & Dhikav, 2012)</u>, is a complex and highly plastic neural structure <u>(Fuchsberger & Paulsen, 2022)</u> that exhibits specialized, lateralized functionality. The left hippocampus is primarily associated with context-specific autobiographical memory, while the right hemisphere is mainly involved in spatially-dependent processes <u>(Burgess et al., 2002; Robinson et al., 2016)</u>. Additionally, the hippocampus also exhibits functional diversity along its long (i.e., anterior-posterior) axis <u>(Strange et al., 2014)</u>, where the anterior portions are associated with novelty detection, imagination, schemata representations, and the recollection of past events (<u>Kafkas & Montaldi, 2018; Robin & Moscovitch, 2017;</u> <u>Zeidman & Maguire, 2016</u>), the intermediate hippocampus is associated with the rapid encoding of spatial information from the external environment <u>(Blanquat et al., 2013)</u> and shares functional characteristics with both anterior and posterior parts <u>(Fanselow & Dong, 2010)</u>, and the posterior parts are primarily associated with sensorimotor and spatial processing <u>(Wang et al., 2022)</u>. Furthermore, the hippocampus contributes to a wide range of cognitive functions related to learning and memory <u>(Maguire, 2014; Wiltgen et al., 2010)</u>. This versatility is due to its robust connections with other brain regions important to numerous higher-order cognitive processes that are utilized during physics learning, ranging from spatial cognition to numerical processing, such as the entorhinal, frontal, and parietal cortices <u>(Anand & Dhikav, 2012; Das & Menon, 2021; A.</u> <u>Zheng et al., 2021)</u>. Moreover, supporting constructivist epistemology, the hippocampus exhibits heightened sensitivity to associative novelty that arises from a mismatch between retrieved memory representations and new sensory inputs. As a result, it plays a critical role in comparing these novel inputs with existing memories and assimilating new information into pre-existing memory structures <u>(Kragel et al., 2020)</u>. Furthermore, a prior study has found that hippocampal connectivity changes with respect to learning methodologies. Specifically, <u>Voss et al., (2011)</u> found that volitional learning, when compared to passive learning, increased hippocampal engagement and its connectivity with regions such as the bilateral dorsolateral prefrontal cortex, and cerebellum, resulting in improved spatial accuracy on subsequent recall tests.

Building on these prior findings, we hypothesized that active learning approaches in physics education promote differential engagement of the hippocampus compared to traditional lecture-based methods. Using functional magnetic resonance imaging (fMRI), we examined hippocampal whole-brain functional connectivity changes across a semester of introductory university physics to determine any differences among physics students enrolled in either modeling-instruction, an active learning approach that emphasizes collaborative problem solving and learning through activities <u>(Brewe et al., 2018)</u>, or traditional lecture-based instruction. To this end, we assessed connectivity changes (i.e., pre-vs. post-instruction) across three cognitive contexts: physics conceptual reasoning, physics knowledge retrieval, and rest.

First, we assessed the general effects of physics education by examining pre-to post-instruction changes in hippocampal whole-brain functional connectivity (i.e., main effect of time). In line with constructivist theory, we anticipated that as students participated in activities and attended lectures about Newtonian-based physics phenomena, their naive conceptualizations would be reduced as they developed more scientifically-founded schemata. Given the role of the hippocampus in both memory updating and relational memory, we anticipated that during physics conceptual reasoning, physics knowledge retrieval, and rest, students would exhibit increased hippocampal connectivity at post-instruction, regardless of instructional methodology. Second, we examined hippocampal connectivity changes associated with pedagogical effects, including modeling vs. lecture-based instruction (i.e., main effect of classroom), as well as the interaction between classroom and time. Consistent with constructivist theory, as modeling instruction students typically engage in more experiential learning, we hypothesized for both the main effect of classroom, as well as the interaction between classroom and time, that modeling instruction students would show enhanced hippocampal functional connectivity across physics conceptual reasoning, physics knowledge retrieval, and rest. Third, for all effects (i.e., time, classroom, and time × classroom), we examined associations between changes in hippocampal connectivity and physics task accuracy. We predicted that increased task-based connectivity would be associated with improved accuracy across time, especially for modeling students. Consistent with constructivist theory, this would reflect a more integrated understanding of physics concepts, as active learning approaches are intended to support the development of more scientifically-grounded mental schemata.

Finally, we performed an exploratory analysis investigating whether hippocampal connectivity changes observed during physics tasks and rest generalized to improved performance on measures of general intellectual ability. We examined associations between hippocampal connectivity patterns during physics conceptual reasoning and cognitive abilities of perceptual reasoning and working memory, as well as during physics knowledge retrieval and verbal comprehension. We also examined associations between hippocampal connectivity during rest and cognitive abilities of perceptual reasoning, working memory, verbal comprehension, and processing speed, as prior findings suggest that resting state can be broadly predictive of cognitive task performance <u>(Cole et al., 2016)</u>. Collectively, we sought to demonstrate that connectivity changes linked to physics instruction may extend beyond physics-related content, suggesting broader cognitive benefits of physics education.

## Materials and Methods

### Participants

The study sample comprised 121 healthy, right-handed undergraduate students (mean age = 19.8 ± 1.5 years; range: 18–26 years; 55 females) enrolled in a 15-week semester of calculus-based introductory physics at Florida International University (FIU). Participants received one of two instructional methods: traditional lecture-based instruction (*n*=60) or modeling instruction (*n*=61), an active learning approach that engages students in model building and collaborative group activities to understand physics concepts. Both classrooms covered topics related to Newtonian mechanics, including motion along straight lines and in two and three dimensions, Newton’s laws of motion, work and energy, momentum and collisions, and rotational dynamics. Upon study enrollment, participants provided information on their age, sex, ethnicity, household income, grade point average (GPA), and academic standing (i.e., freshman, sophomore, junior, or senior) (**Table 1**). All participants were free from cognitive, neurological, or psychiatric conditions and reported no use of psychotropic medications. Furthermore, to mitigate concerns about variability in academic ability and performance, all participants had either a GPA greater than 2.24 or an SAT math score greater than 500.

**Table 1.** Participant Information.

|  | Modeling Instruction (n=61) |  | Lecture-Based Instruction (n=60) |  |
| --- | --- | --- | --- | --- |
|  | N | Percentage | N | Percentage |
| **Gender** |  |  |  |  |
| Male | 37 | 61 | 31 | 52 |
| Female | 24 | 39 | 29 | 48 |
| **Ethnicity** |  |  |  |  |
| Hispanic | 44 | 72 | 43 | 72 |
| Non-Hispanic | 17 | 28 | 17 | 28 |
| **Household Income** |  |  |  |  |
| < \$15,000 | 12 | 20 | 13 | 22 |
| \$15,000 - \$34,999 | 10 | 16 | 16 | 27 |
| \$35,000 - \$49,999 | 6 | 10 | 9 | 15 |
| \$50,000 - \$74,999 | 15 | 25 | 7 | 12 |
| \$75,000 - \$99,999 | 11 | 18 | 6 | 10 |
| >\$100,000 | 7 | 11 | 9 | 15 |
| **Years Enrolled** |  |  |  |  |
| Freshman | 6 | 10 | 6 | 10 |
| Sophomore | 27 | 44 | 29 | 48 |
| Junior | 21 | 34 | 18 | 30 |
| Senior | 7 | 11 | 7 | 12 |
|  | **Grand Mean** | **Std. Dev.** | **Grand Mean** | **Std. Dev.** |
| **Age** | 19.84 | 1.67 | 19.73 | 1.43 |
| **GPA** | 3.29 | 0.67 | 3.29 | 0.38 |

Note. “N” represents the sample size of the group, and “Percentage” represents the percentage of participants in that group for each categorical variable (i.e., gender, ethnicity, household income, and years enrolled).

### Procedures

Behavioral and imaging data collection occurred during two time periods: pre- and post-instruction. At the beginning of the academic semester, different recruitment strategies were employed, including in-class visits, emailing eligible students, and distributing flyers across campus. Enrolled participants completed an fMRI session at the start of the course (i.e., pre-instruction), no later than the fourth week of instruction and before the first exam. Imaging sessions were conducted off-campus, with participants provided free parking and/or transportation to and from the MRI site. At the end of the semester, participants completed their post-instruction session after the final exam but no later than two weeks after the semester concluded. Written informed consent was obtained in accordance with FIU’s Institutional Review Board approval. Additionally, participants were monetarily compensated for their study participation.

### Measures of Cognitive Abilities

Participants completed the Wechsler Adult Intelligence Scale-Fourth Edition (WAIS-IV; <u>Wechsler, 2012</u>) at both the pre- and post-instruction sessions. The WAIS-IV provides a comprehensive assessment of adult cognitive abilities across four domains. The verbal comprehension index (VCI) measures lexical knowledge, semantic understanding, and verbal reasoning of conceptually related words. The perceptual reasoning index (PRI) assesses non-verbal abstract problem solving through spatial reasoning tasks involving pattern recognition, component-based puzzle reconstruction, and translation of three-dimensional designs into two-dimensional representations. The working memory index (WMI) evaluates short-term memory capacity using auditory stimuli and mental arithmetic. Lastly, the processing speed index (PSI) measures information processing efficiency through tests requiring rapid visual scanning to identify specific targets among irrelevant stimuli and visuomotor coordination to record symbol-number associations <u>(Hartman, 2009)</u>.

### MRI Data Acquisition

MRI data were acquired on a GE 3T Healthcare Discovery 750W MRI scanner at the University of Miami. Functional imaging data were acquired with an interleaved gradient-echo, echo planar imaging (EPI) sequence (repetition time [TR]/echo time [TE] = 2000/30ms, flip angle = 75°, field of view (FOV) = 220x220mm, matrix size = 64x64, voxels dimensions = 3.4×3.4×3.4mm, 42 axial oblique slices). T1-weighted structural data were also acquired using a 3D fast spoiled gradient recall brain volume (FSPGR BRAVO) sequence with 186 contiguous sagittal slices (TI = 650ms, bandwidth = 25.0kHz, flip angle = 12°, FOV = 256x256mm, and slice thickness = 1.0mm).

### fMRI Tasks

#### Physics Conceptual Reasoning Task

Participants completed a self-paced, conceptual reasoning task consisting of questions derived from the Force Concept Inventory (FCI) (**Fig. 1A-C**) (<u>Hestenes et al., 1992; Lasry et al., 2011; Von Korff et al., 2016</u>) that was adapted for the MRI environment <u>(Bartley et al., 2019)</u>. FCI trials included textual descriptions and illustrations of scenarios, including objects at rest or in motion, accompanied by one correct Newtonian solution and several incorrect non-Newtonian solutions. Control trials included similar visual stimuli to FCI trials but tested general reading comprehension and/or shape discrimination. Each trial was presented in blocks composed of three sequential view screens: Phase I: Scenario, which included an illustration and text describing a physical scenario; Phase II: Question, which involved presenting a question related to the scenario; and Phase III: Answer, wherein participants were presented with four answer choices. Both FCI and control blocks had a maximum duration of 45 sec and were followed by a fixation cross with a minimum duration of 10 sec. The total duration for each FCI run was 5 min 44 sec; data were collected during three runs for a total duration of ∼16 minutes, resulting in 170 volumes being collected per participant.

**Figure 1.**
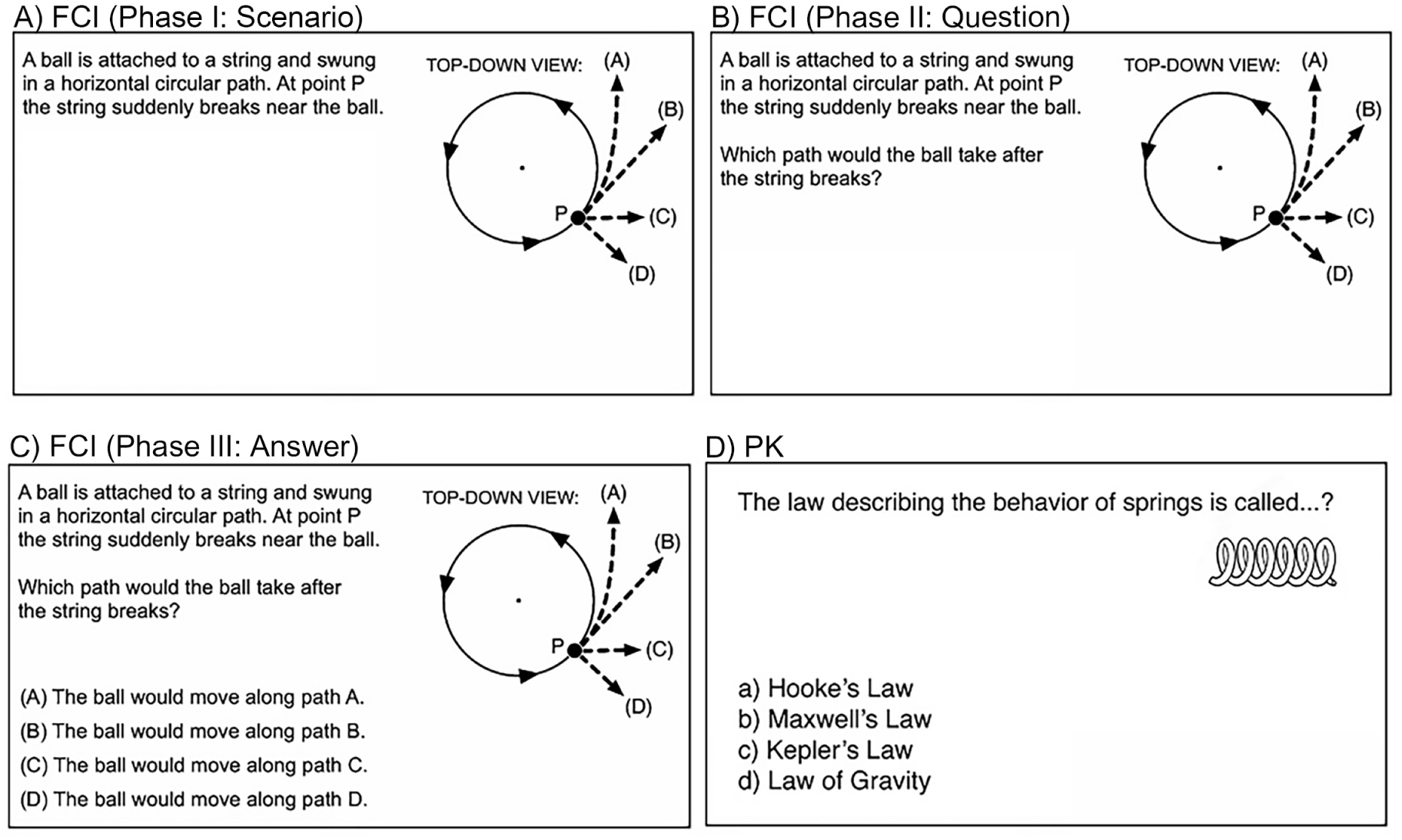
Force Concept Inventory (FCI) and Physics Knowledge (PK) Tasks. Example items of the in-scanner physics tasks, including the three phases of the FCI task (A) Phase I: Scenario, (B) Phase II: Question, (C) Phase III: Answer, and (D) the PK task. Reprinted from Trends in Neuroscience and Education, Vol 32, <u>Smith et al., 2023</u>, Task-based attentional and default mode connectivity associated with science and math anxiety profiles among university physics students, Pages 100204, Copyright (2023), with permission from Elsevier.

#### Physics Knowledge Retrieval Task

Participants also completed the physics knowledge (PK) task (**Fig. 1D**), which was adapted from a general knowledge task of semantic retrieval <u>(Elman et al., 2012)</u>. The PK task was presented in a block design and probed for brain activation associated with physics-based content knowledge. Students viewed physics questions related to general physics knowledge and their corresponding answer choices. A control condition was presented in which students viewed general knowledge questions with corresponding answer choices. PK and control blocks were 28 seconds long and included four questions per block (6.5 sec per question followed by 0.5 sec of quick fixation). Three blocks of physics or general questions (six question blocks total) were alternated with 10 sec of fixation. The total duration of one run was 4 min 2 sec; data were collected during two runs for a total duration of ∼8 minutes, resulting in 180 volumes being collected per participant.

#### Resting State

Participants were instructed to lie quietly with their eyes closed and not to fall asleep. The total duration of the resting state session was 12 minutes, resulting in 365 volumes of data being collected per participant. The first five volumes for each participant were removed to ensure only steady-state data were used in subsequent analyses, leaving 360 volumes of resting state data per participant.

### Analyses

#### fMRI Preprocessing

As stated in <u>Smith et al., 2023</u>, the functional runs for all tasks/states used within this study (i.e., FCI, PK, rest) were preprocessed independently. Each participant’s T1-weighted images were corrected for intensity non-uniformity with `ANT’s N4BiasFieldCorrection` tool <u>(Avants et al., 2008; Tustison et al., 2010)</u>. Anatomical and functional images underwent additional preprocessing steps using `fMRIPrep` (v.1.5.0rc1) <u>(Esteban et al., 2020)</u>. The T1-weighted (T1w) reference, which was used throughout the pipeline, was generated after T1w images were corrected for intensity non-uniformity with `ANT’s N4BiasFieldCorrection`. `Freesurfer`’s `mri_robust_template` was used to generate a T1w reference, which was used throughout the entire pipeline <u>(Reuter et al., 2010; Tustison et al., 2010)</u>. `Nipype’s` implementation of `ANT’s antsBrainExtraction` workflow was used to skullstrip the T1w reference using `OASIS30ANTs` as the target template (<u>Gorgolewski et al., 2011</u>). `FSL’s FAST` was used for brain tissue segmentation of the cerebrospinal fluid (CSF), white matter (WM), and gray matter (GM); brain surfaces were reconstructed using `Freesurfer’s recon_all` (Dale et al., 1999; Zhang et al., 2001). Preprocessing of functional images began with selecting a reference volume and generating a skullstripped version using a custom methodology of `fMRIPrep`. `Freesurfer’s bbregister`, which uses boundary-based registration, was used to coregister the T1w reference to the BOLD reference. The BOLD time series was then resampled onto surfaces of `fsaverage5` space and resampled onto their original, native space by applying a single, composite transform to correct for head motion and susceptibility distortions. Additionally, the BOLD time series was high pass filtered, using a discrete cosine filter with a cutoff of 128s <u>(Greve & Fischl, 2009)</u>. Several confounding time series were estimated as follows: for each functional run, motion outliers were set at a threshold of 0.5 mm framewise displacement (FD). Nuisance signals from the CSF, WM, and whole brain masks were extracted by using a set of physiological regressors, which were extracted to allow for both temporal component-based noise correction (tCompCor) and anatomical component-based noise correction (aCompCor) <u>(Behzadi et al., 2007)</u>. Additionally, the confound time series derived from head motion estimates were expanded to include its temporal derivatives and quadratic terms, resulting in a total of 24 head motion parameters (i.e., six base motion parameters, six temporal derivatives of six motion parameters, 12 quadratic terms of six motion parameters, and their six temporal derivatives). Estimates for the global, cerebrospinal fluid, and white matter signals were expanded to include their temporal derivatives and quadratic terms, resulting in a total of 12 signal-based parameters (i.e., three base signal parameters, three temporal derivatives of the three base parameters, the three quadratic terms of the base parameters, and the three quadratic terms of the temporal derivatives). Finally, all 24 head motion confound estimates, three high pass filter estimates, and a variable number of aCompCor estimates (components that explain the top 50% of the variance) were outputted into a tsv file to be used for later denoising steps <u>(Satterthwaite et al., 2013)</u>. An expanded description of the preprocessing workflow can be found in the Supplementary Material (SM).

#### Quality Control Assessment

`MRIQC` <u>(Esteban et al., 2017)</u> was used to compute quality control metrics for each participant’s functional runs, including mean framewise displacement, ghost-to-signal ratio, signal-to-noise ratio, and entropy focus criterion, which quantifies head-motion related artifacts such as blurring and ghosting <u>(Atkinson et al., 1997)</u>. Participants’ functional runs with a mean framewise displacement, ghost-to-signal ratio, or entropy focus criterion above the 99th percentile, or a signal-to-noise ratio below the 1st percentile, were excluded from the analysis. The application of this quality control procedure led to the exclusion of complete functional run datasets for several participants across different experimental conditions. For resting state data, two participants had all functional runs excluded from both pre- and post-instruction sessions. Similarly, complete datasets were excluded for two participants in the FCI task and one participant in the PK task, with these exclusions overlapping with the resting state exclusions. Additional exclusions occurred at the session level where twelve participants had all functional runs excluded from either the pre-instruction (n=6) or post-instruction (n=6) resting state session. Overall, MRIQC identified fourteen unique participants with incomplete datasets for at least one experimental condition (resting state, FCI, or PK). These exclusions were primarily attributed to excessive motion artifacts (including blurring and ghosting) and inadequate signal-to-noise ratio (e.g., thermal noise).

### Denoising and Time Series Extraction

All fMRI data were subjected to the same denoising and extraction strategies. Analyses were restricted to subjects that possessed at least a single functional run for each task and session, resulting in 90 participants included in the analyses. Thus, any potential differences were not confounded by tasks utilizing different subject pools, enhancing overall interpretability. Additional participant demographic information for these participants can be found in the SM.

For each run of fMRI data, a design matrix comprising the first six anatomical components derived from the white-matter and cerebrospinal fluid (CSF) mask, the six head motion parameters and their first derivatives, and the global signal and its first derivative was constructed. Nuisance regression was employed using `AFNI’s 3dtproject` to orthogonalize each run to each confound of interest. Furthermore, each run was also band-pass filtered (0.008-0.1 Hz), spatially blurred (fwhm=6mm) to enhance the signal to noise ratio, and standardized. After nuisance regression, the task-based volumes corresponding to the physics conditions of each task were isolated and retained using `AFNI’s 3dTcat` (i.e., control trials were removed), resulting in approximately 40 and 45 frames retained per run for the FCI and PK tasks, respectively. To account for the hemodynamic delay <u>(Buxton et al., 2004)</u>, onset times for each physics trial block were shifted forward by 4 seconds <u>(Wittkuhn & Schuck, 2021)</u>. Additionally, volumes exceeding a framewise displacement of 0.35mm were scrubbed using `3dcat`. Furthermore, each subject was assessed to ensure that the proportion of volumes removed did not exceed 40% per run for resting state data and 20% per run for the physics-only conditions of the task data (i.e., FCI, PK). A stricter retention threshold was applied to the task data due to the difference in the number of frames being assessed (i.e., 360 frames for rest compared to 40 and 45 frames per run for the FCI and PK tasks, respectively). Once censoring was completed, each run from the FCI and PK tasks were concatenated into a single, continuous run using `3dTcat`, such that the FCI and PK task were treated as a single run for the subsequent connectivity analysis.

#### Connectivity Analyses

After denoising concatenation, participant-specific first-level functional connectivity analyses were performed. First, the hippocampus was parcellated into six seed regions (i.e., three per hemisphere) using a meta-analytically-derived hippocampal atlas <u>Plachti et al., 2019</u> (**Fig.2**) and resampled to *MNI152NLin2009cAsym* space using `AFNI’s 3dresample`. For each seed, time series data were extracted using AFNI’s `3dmaskave`, averaged to create a single time series per seed, and incorporated into a design matrix. Separate design matrices were constructed for each hemisphere, ensuring that seed time series within the same hemisphere accounted for each other’s variability. Subsequently, for each participant, generalized least squares (GLS) regression was conducted using `AFNI’s 3dREMLfit` to compute beta weights between each hippocampal seed and non-ROI voxels, while accounting for temporal autocorrelation within the data.

**Figure 2.**
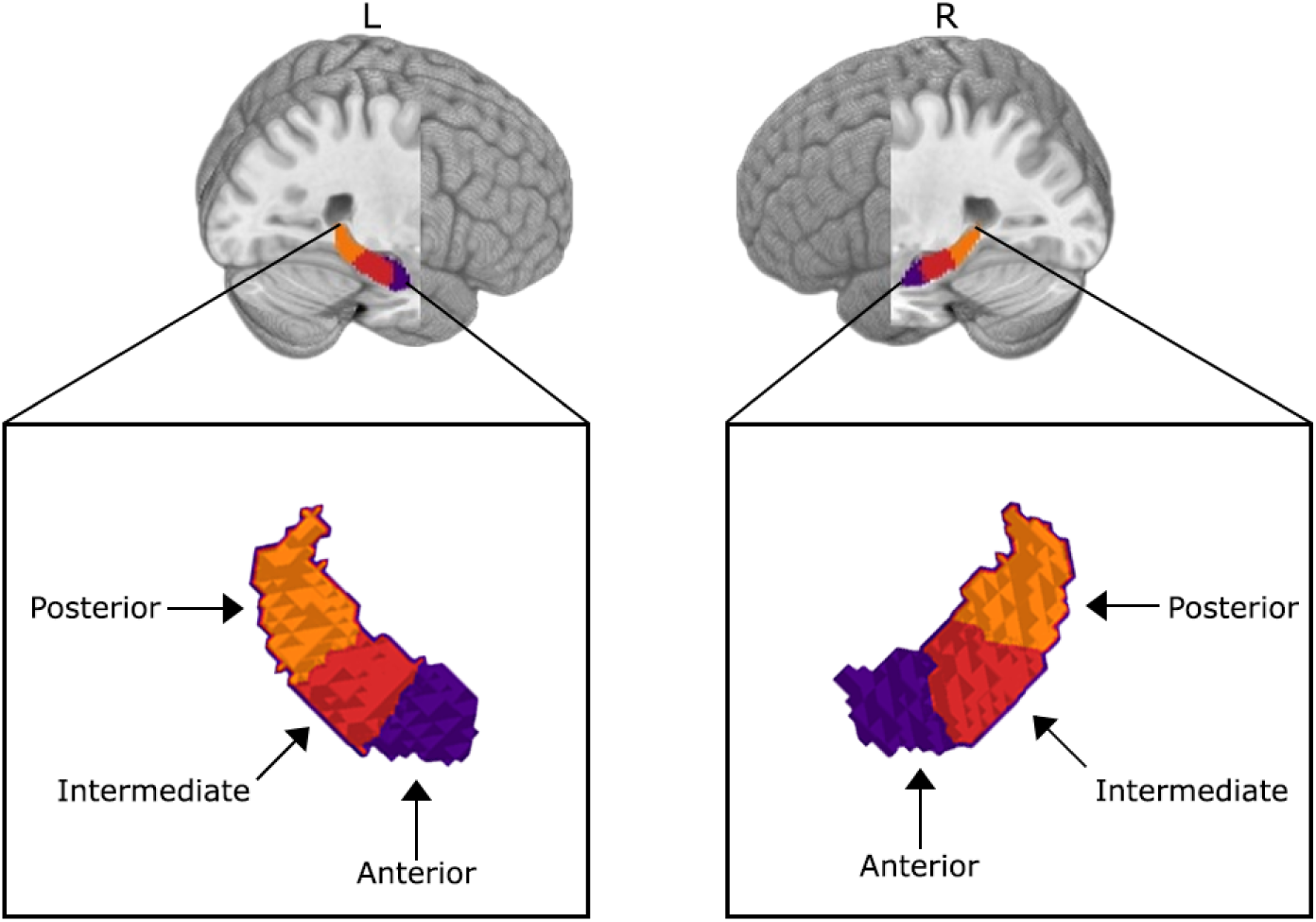
Hippocampal Parcellation. In the current study, a meta-analytically derived hippocampal parcellation was utilized <u>Plachti et al., 2019</u>). For each participant, the left and right hippocampus were subdivided into three seeds (resulting in six seeds total), including the anterior (purple), intermediate (red), and posterior (orange) hippocampus.

Following the first-level analysis, the resulting beta weights were utilized in the group-level analysis. A sample-specific group mask was created using `Nilearn’s intersect_mask` function, retaining only those voxels where at least 50% of participants had non-zero values. A linear mixed-effects regression analysis was performed using `AFNI’s 3dLMEr`, generating contrasts for the main effects of time (i.e., pre-vs. post-instruction), classroom (i.e., modeling vs. lecture-based instruction), and their interaction (i.e., time × classroom). This model also controlled for nuisance variables, including age, sex, ethnicity, income, years of education, and subject-specific random effects. Additionally, the regression produced three group level contrast maps (z-maps) corresponding to each contrast of interest and a BRIK file containing the residuals.

To estimate spatial smoothing parameters, the residual BRIK file was processed with `AFNI’s 3dFWHMx`, which employs an autocorrelation method that does not assume Gaussian noise. These parameters were then input to `AFNI’s 3dClustSim` to determine the appropriate cluster size for controlling the family-wise error rate (FWE) at several predefined voxel thresholds, which were used to assess each group-level contrast map.

#### Statistical Analyses

First, we examined the extent to which modeling and lecture-based instruction students differed with respect to age, sex, ethnicity, household income, number of years enrolled at FIU (i.e., freshman, sophomore, junior, or senior), and GPA (both pre- and post-instruction). To this end, *t*-tests and *chi-square* tests were used to test the entire sample (𝑛=121), as well as the subsample of participants with at least one functional run of all tasks and sessions (𝑛=90).

Second, group-level connectivity analyses examined hippocampal connectivity differences across each contrast, which included the main effects of time (i.e., pre-vs. post-instruction), classroom (i.e., modeling vs. lecture-based instruction), and their interaction (i.e., time × class). Each group-level contrast map was thresholded using `Nilearn’s threshold_img` function to retain only voxels with z-scores corresponding to 𝑝*_voxel-wise_*<0.001. Significant clusters were identified using `Nilearn’s get_cluster_table`, which uses the NN1 definition, where voxels are considered part of the same cluster only if their faces touch. Only clusters meeting a significance threshold of 𝑝*_FWE-corrected_*<0.05 were retained. The MNI coordinates within significant clusters, which defined the center of mass of each cluster, were labeled using the Eickhoff-Zilles macro labels atlas from `AFNI’s whereami` program.

Finally, for each significant cluster identified from the group-level connectivity analyses, binary masks for these clusters were generated. These binary masks were used to extract the averaged, non-zero beta weights from the individual, first-level analysis maps for the exploratory behavioral analyses. For the FCI task, a linear regression model was used to assess the association between the change (i.e., post-vs. pre-instruction) in (i) FCI accuracy, (ii) PRI, and (iii) WMI and the change in beta coefficients for the (i) main effect of time, (ii) main effect of class, (iii) and the interaction between time and classroom. For the PK task, a linear regression model was used to assess the association between the change in (i) PK accuracy, (ii) WMI, and (iii) VCI) and the change in beta coefficients for the (i) main effect of time, (ii) main effect of class, (iii) and the interaction between time and classroom. Finally, for rest, a linear regression model was used to assess the association between the change in (i) PRI, (ii) WMI, (iii) VCI, and (iv) PSI and the change in beta coefficients for the (i) main effect of time, (ii) main effect of classroom, (iii) and the interaction between time and classroom.. All regression models controlled for age, sex, ethnicity, household income, years enrolled at FIU, and GPA.

#### Functional Decoding of Statistical Maps

For all contrasts (main effect of time, main effect of class, and interaction effect between time and class), each thresholded z-map exhibiting significant clusters after correction was used to perform a functional decoding analysis to determine the functional role of each significant cluster. To this end, we used NiCLIP <u>(Peraza et al.,</u> <u>2025)</u>, a neuroimaging contrastive language-image pretraining framework that predicts text from brain images using a CLIP architecture with text encoding enhanced with BrainGPT <u>(Luo et al., 2024)</u>, a pre-trained Large-Language Model (LLM) fine-tuned on neuroscience papers. NiCLIP was trained on text and brain activation coordinates from approximately 24,000 fMRI articles obtained through PubMed Central. Text embeddings were generated using BrainGPT, while image embeddings were created by first calculating MKDA-modeled activation brain maps from peak coordinates by convolving with a binary sphere of fixed radius, then applying spatial dimensionality reduction using the DiFuMo atlas. For our decoding analysis, we used this pre-trained CLIP model and the braindec package (https://github.com/NBCLab/brain-decoder) <u>(Peraza et al., 2025)</u> to encode the task names and definitions of the Cognitive Atlas <u>(Poldrack et al., 2011)</u>, an online database of concepts in cognitive science, into text embeddings. We encoded our statistical maps into image embeddings using the same DiFuMo parcellation. The posterior probability of predicted tasks was computed based on the cosine similarity between text and image embeddings in the shared latent space, which were then converted to probabilities using the softmax function. Finally, these posterior probabilities were used to create visualizations of the top five concepts for each statistical map demonstrating significant functional connectivity with the hippocampus.

## Results

### Participant Differences Between Active Learning and Lecture-Based Classrooms

No significant differences were observed between students in the active learning classrooms (𝑛=61) and those in the lecture-based classrooms (𝑛=60) in terms of age, sex, ethnicity, household income, years enrolled at FIU, and GPA (**Supplementary Table 2**). Among the subset of participants with at least one functional run for all tasks and sessions, which included active learning (𝑛=46) and lecture-based students (𝑛=44), we also observed no significant differences across participant groups (**Supplementary Table 2**).

### Hippocampal Whole-Brain Connectivity

We observed differences in hippocampal connectivity across time (i.e., main effect of time), classroom (i.e., main effect of class), and for the interaction between time and classroom (**Table 2**).

**Table 2.** Connectivity Analyses: Main Effect of Time, Main Effect of Classroom, and Interaction Effects. Hippocampal whole-brain connectivity analyses were conducted for each hippocampal seed region (i.e., anterior, intermediate, posterior) for each hemisphere (i.e., left, right). Significant clusters (𝑝_𝐹𝑊𝐸−𝑐𝑜𝑟𝑟𝑒𝑐𝑡𝑒𝑑_<0.05) are shown for the main effect of time (i.e., pre- > post-instruction) and the interaction effect (i.e., modeling > lecture-based, lecture-based > modeling) across the FCI Task, PK Task, and Rest. No significant effects were observed for increasing main effects of time (i.e., post- > pre-instruction) or for the main effects of the classroom. For each cluster, the peak MNI coordinate, along with the cluster extent (𝑚𝑚^3^), z-value, and the corresponding anatomical label are provided.

| Hippocampal Seed | MNI Coordinate |  |  | Cluster Extent (mm<sup>3</sup>) | Z-value | Region Label |
| --- | --- | --- | --- | --- | --- | --- |
|  | X | Y | Z |  |  |  |
| Main Effect of Time |  |  |  |  |  |  |
| Pre-Instruction > Post-Instruction |  |  |  |  |  |  |
| FCI Task |  |  |  |  |  |  |
| L. Intermediate | 41.5 | 15.5 | 27.5 | 2792 | -4.97 | R. Inferior Frontal Gyrus (Opercularis) |
|  | 49.5 | 53.5 | 5.5 | 2112 | -4.73 | R. Middle Frontal Gyrus |
| Rest |  |  |  |  |  |  |
| L. Intermediate | -4.5 | -52.5 | 23.5 | 3944 | -5.05 | L. Precuneus |
|  | -4.5 | -46.5 | 71.5 | 3256 | -5.03 | L. Precuneus |
| R. Anterior | -14.5 | -52.5 | -0.5 | 1200 | -4.90 | L. Lingual Gyrus |
|  | -50.5 | -18.5 | 53.5 | 2024 | -4.72 | L. Postcentral Gyrus |
|  | 17.5 | -74.5 | -0.5 | 1856 | -4.55 | R. Lingual Gyrus |
| Pre-Instruction < Post-Instruction |  |  |  |  |  |  |
| n.s. |  |  |  |  |  |  |
| Main Effect of Instruction |  |  |  |  |  |  |
| Modeling-Instruction > Lecture-Based Instruction |  |  |  |  |  |  |
| n.s. |  |  |  |  |  |  |
| Modeling-Instruction < Lecture-Based Instruction |  |  |  |  |  |  |
| n.s. |  |  |  |  |  |  |
| Interaction: Time X Instruction |  |  |  |  |  |  |
| Modeling-Instruction > Lecture-Based Instruction |  |  |  |  |  |  |
| FCI Task |  |  |  |  |  |  |
| R. Anterior | -30.5 | -60.5 | 49.5 | 1272 | 4.67 | L. Superior Parietal Lobule |
| PK Task |  |  |  |  |  |  |
| L. Anterior | -34.5 | -74.5 | -36.5 | 1104 | 5.09 | L. Cerebellum (Crus 2) |
|  | 43.5 | -70.5 | -22.5 | 1176 | 4.50 | R. Cerebellum (Crus 1) |
| Rest |  |  |  |  |  |  |
| L. Intermediate | -12.5 | 49.5 | 31.5 | 1264 | 5.08 | L. Superior Frontal Gyrus |
| Modeling-Instruction < Lecture-Based Instruction |  |  |  |  |  |  |
| Rest |  |  |  |  |  |  |
| L. Intermediate | 31.5 | -40.5 | -12.5 | 2520 | -5.64 | R. Fusiform Gyrus |

### Main Effect of Time: Pre-vs. Post-Instruction

During the FCI task, significantly decreased connectivity was observed at post-compared to pre-instruction for the left intermediate hippocampus and the right inferior and middle frontal gyri (**Fig. 3A**). The decoding analysis suggested that the top concepts associated with this pattern were *visual object recognition*, *visual perception*, *autobiographical memory*, and self-monitoring. During rest, decreased post- to pre-instruction connectivity was observed between the left intermediate hippocampus and the left precuneus (**Fig. 3B**), a functional pattern that the decoding analysis revealed was associated with *visual perception*, *visual object recognition*, *social cognition*, *autobiographical memory*, and *self-monitoring*. Additionally, during rest, decreased post- to pre-instruction connectivity was observed between right anterior hippocampus and left postcentral gyrus and bilateral lingual gyri (**Fig. 3B**), a functional pattern where the top decoded concepts were *autobiographical memory*, *self-monitoring*, *visual object recognition*, and *visual perception*. Significant decreases in hippocampal connectivity during the PK task were not observed. No significant increases in hippocampal connectivity were observed post-compared to pre-instruction.

**Figure 3.**
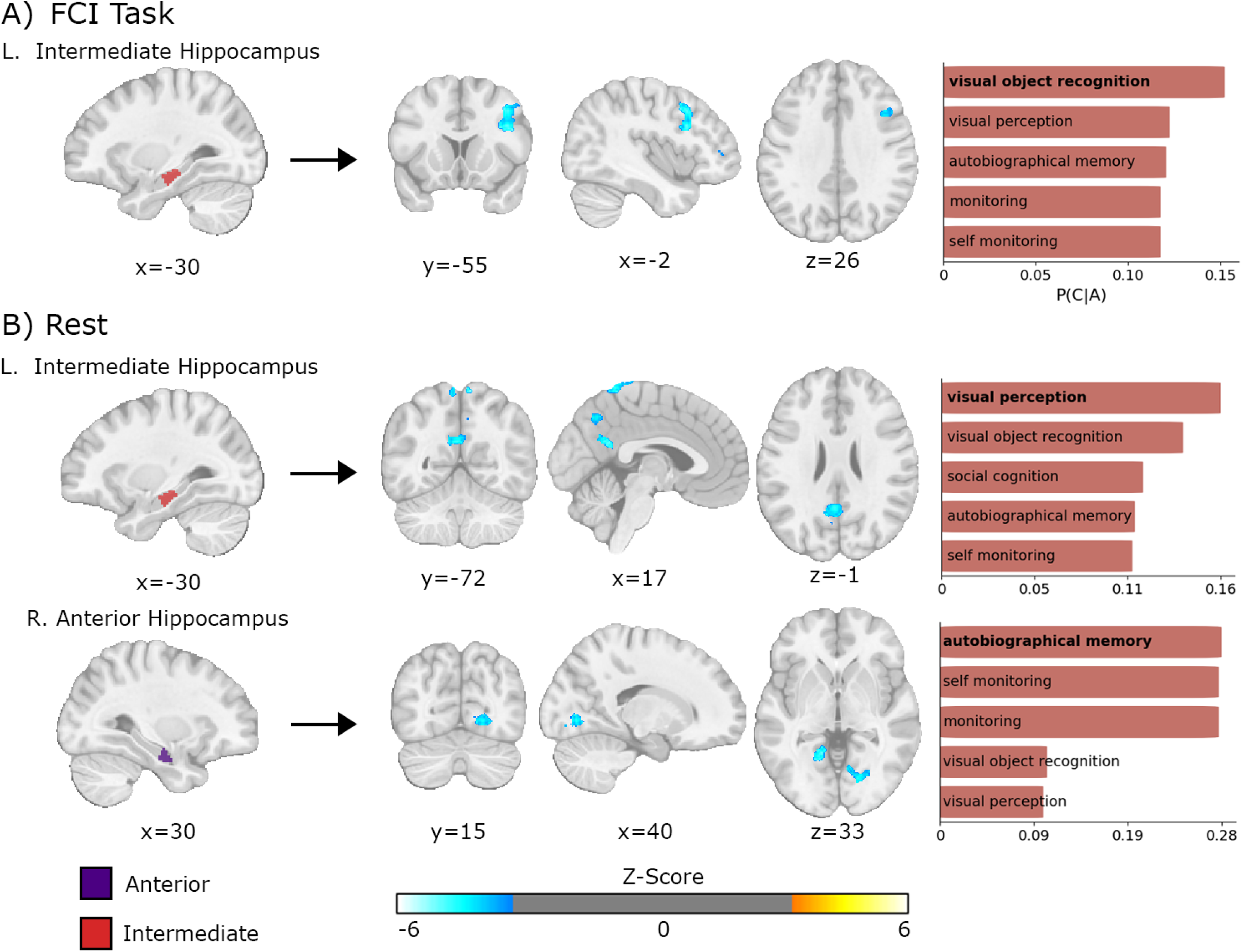
Main Effect of Time. Significant decreases in hippocampal connectivity at post-compared to pre-instruction were observed during (a) the FCI Task and (b) Rest. Seed regions for the intermediate and anterior hippocampus are shown in red and purple, respectively, alongside resultant connectivity maps in which negative *z*-values (blue) indicate decreased connectivity at post-relative to pre-instruction. Decoding results in which the top the concepts with the top five highest probabilities associated given the activation pattern - P(C|A), the probability of a particular psychological concept given the pattern of activation in the statistical map - are depicted by the graph on the right.

### Main Effect of Classroom: Active Learning vs. Lecture-Based Instruction

No significant results were observed for the main effect of classroom (i.e., Modeling > Lecture-Based or Lecture-Based > Modeling) for the FCI or PK tasks or rest.

### Interaction Between Time and Classroom

Results of the interaction analysis revealed four significant clusters across the left and right anterior hippocampus and the left intermediate hippocampus (**Table 2**, **Fig. 4**). Compared to lecture-based students, modeling instruction students exhibited greater changes in connectivity (i.e., post- > pre-instruction) during the FCI task within right anterior hippocampus and left superior parietal lobule, in which the top five decoding results suggests the top concepts are *visual object recognition*, *response selection*, *auditory sentence recognition*, *emotion*, and *reading*, while during the PK task, greater changes in connectivity were observed between left anterior hippocampus and bilateral cerebellum, where the top concepts for the left cerebellum were *response selection*, *emotion*, *response execution*, *feature comparison*, and *reinforcement learning*, while for the right cerebellum the top concepts where *response selection*, *emotion*, *feature comparison*, *visual object recognition*, and *introspection*. During rest, increased connectivity was identified among modeling students relative to lecture-based students within the left intermediate hippocampus and left superior frontal gyrus, with the decoding analysis showing that the top concepts were *visual recognition*, *association learning*, *instrumental condition*, *visual perception*, and *visual object recognition*. Conversely, lecture-based students demonstrated greater changes in connectivity compared to modeling students during rest within the left intermediate hippocampus and right fusiform gyrus, with the decoding results demonstrating that the top concepts were *response selection*, *visual recognition*, *working memory*, *visual working memory*, and *emotion*.

**Figure 4.**
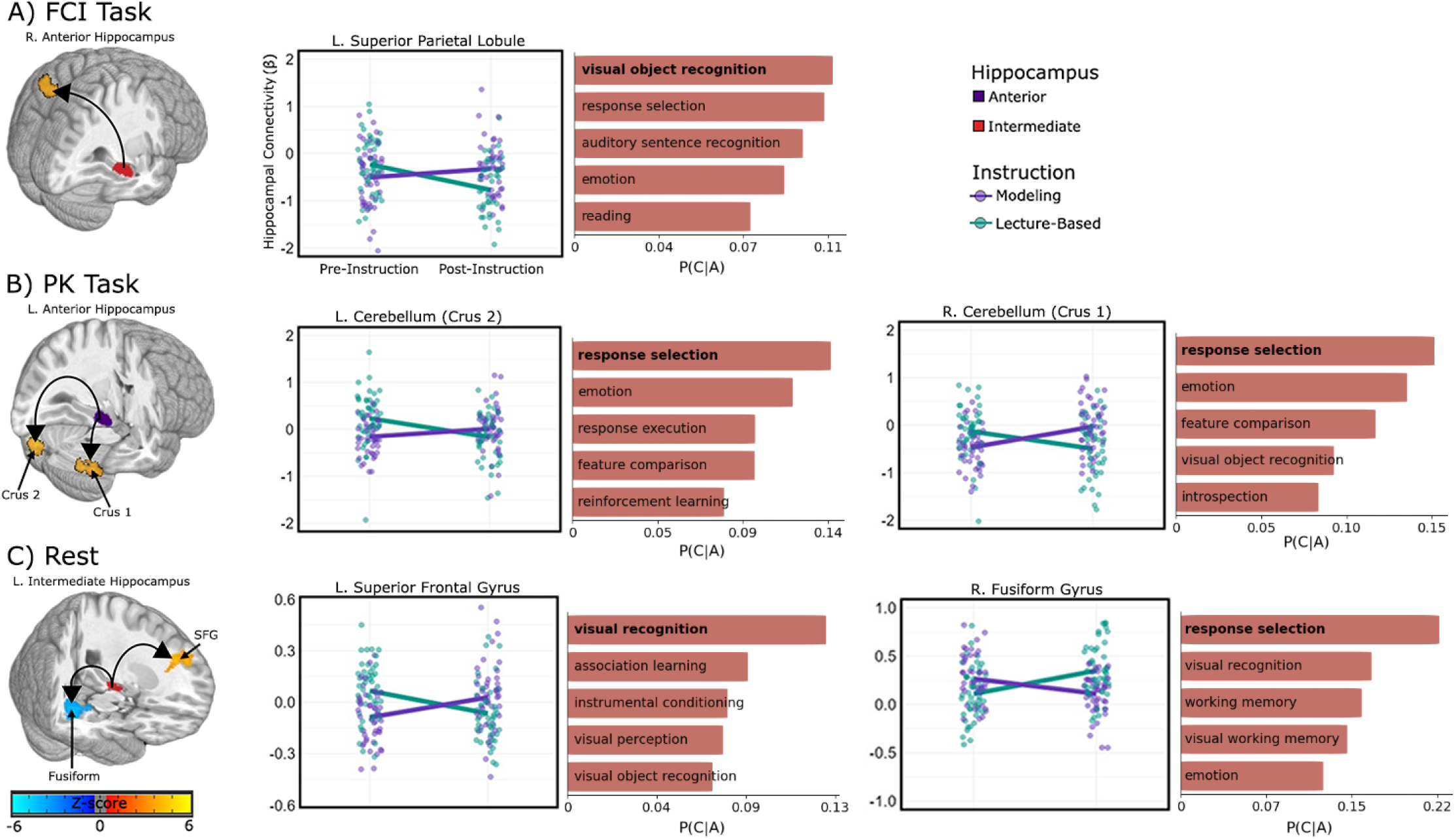
Interaction Effects Between Time and Classroom. Brain renderings display intermediate (red) and anterior (purple) hippocampal seeds and regional clusters for which a significant interaction effect was observed during the (a) FCI Task, (b) PK Task, and (c) Rest. Yellow clusters indicate increasing changes in connectivity (i.e., post- > pre-instruction) among modeling instruction students relative to lecture-based students, whereas blue clusters indicate decreasing changes in connectivity (i.e., pre- > post-instruction). Interaction plots display the corresponding beta coefficients (β) for these regions, reflecting changes in hippocampal connectivity from pre- to post-instruction among modeling instruction (purple) and lecture-based (teal) students. Decoding results in which the top the concepts with the top five highest probabilities associated given the activation pattern - P(C|A) - are depicted by the graph on the right.

### Exploratory Behavioral Effects

The exploratory analyses revealed two significant associations: one between changes in connectivity and changes in task accuracy, and another between changes in connectivity and changes in working memory (Fig. 5). First, increasing changes in connectivity (i.e., post > pre-instruction) between left anterior hippocampus and left cerebellum (crus 2) were associated with increasing PK accuracy among modeling instruction students and decreasing PK accuracy among lecture-based students (β = 0. 0933, 𝑝 = 0. 0259). Second, increasing changes in connectivity between left intermediate hippocampus and right fusiform gyrus were associated with increasing WMI scores among modeling instruction students and decreasing WMI scores among lecture-based students during rest (β = 20. 73, 𝑝 = 0. 0195). All findings for the exploratory behavioral analyses are reported in Supplementary Table 3.

**Figure 5.**
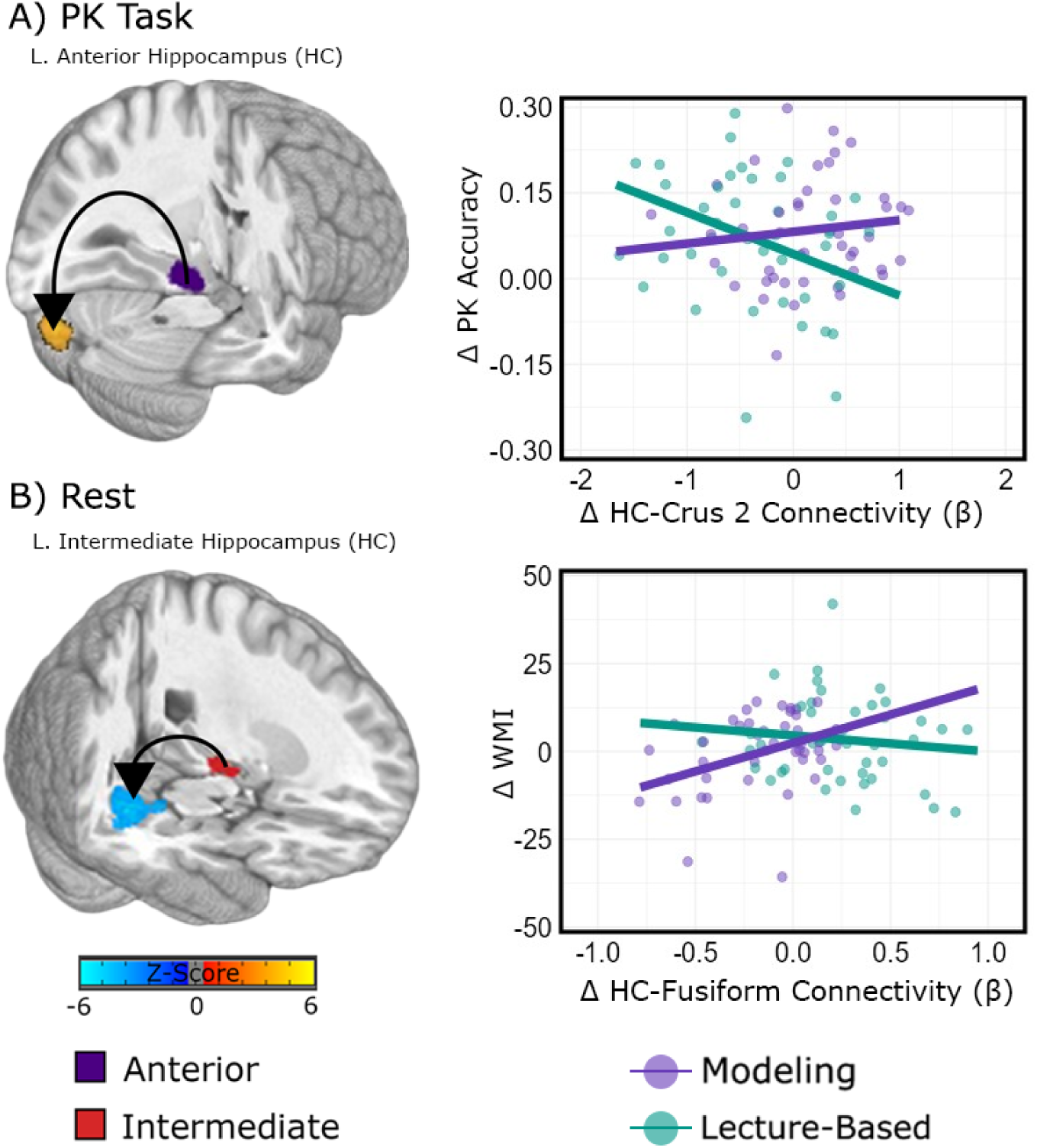
Exploratory Behavioral Effects. Brain renderings display anterior (purple) and intermediate (red) hippocampal seeds and regional clusters for which a significant behavioral effect was observed during the (a) PK Task and (b) Rest. Yellow clusters indicate increasing changes in connectivity (i.e., post- > pre-instruction) among modeling instruction students relative to lecture-based students, whereas blue clusters indicate decreasing changes in connectivity (i.e., pre- > post-instruction). Plots display hippocampal connectivity changes from pre- to post-instruction (Δβ) as a function of (a) changes in PK accuracy and (b) changes in WAIS-IV working memory index (WMI) among modeling instruction (purple) and lecture-based (teal) students.

## Discussion

We investigated differences in hippocampal functional connectivity among university students who completed an introductory physics course taught in either an active learning (i.e., modeling instruction) or lecture-based classroom and examined connectivity changes across time and instructional approach. For the main effect of time, hippocampal connectivity was observed to decrease, not increase as hypothesized, from pre- to post-instruction during the FCI task and rest, with no significant findings observed for the PK task. No effects were observed for the main effect of the classroom (i.e., active learning vs. lecture-based). Findings from the interaction between time and classroom revealed that modeling students exhibited more robust and diverse increases in hippocampal connectivity with parietal, bilateral cerebellar, and frontal regions at post-instruction compared to lecture-based students across all paradigms (i.e., FCI, PK, and rest). In contrast, lecture-based students exhibited increased connectivity with the fusiform gyrus at rest. Finally, during the PK task, task accuracy was predicted by pre- to post instruction hippocampal-cerebellar connectivity changes; similarly during rest, pre- to post-instruction hippocampal-fusiform connectivity changes were predictive of working memory index, but not perceptual reasoning, verbal comprehension, or processing speed. No behavioral associations were observed for the FCI task.

### Pre- to Post-Instruction Changes in Hippocampal Functional Connectivity

During the FCI task, we observed decreased connectivity between the left intermediate hippocampus and both the right inferior frontal gyrus (opercularis) and the right middle frontal gyrus. The left hippocampus is associated with binding together contextual and autobiographical information from retrieved memory traces <u>(Burgess et al., 2002; Robinson et al., 2016)</u>. While the intermediate hippocampus shares functionality with both the anterior and posterior hippocampus, it also has been associated with processing environmental spatial information <u>(Blanquat et al., 2013; Fanselow & Dong, 2010)</u>, suggesting a role in encoding and retrieving relevant contextual information related to spatial information. Each question of the FCI task presents several answer choices related to possible object trajectories, requiring participants to suppress naive responses that may appear intuitively correct but are not scientifically principled. Our findings revealed decreased functional connectivity between the left intermediate hippocampus and two brain regions associated with response inhibition, specifically the right inferior frontal gyrus (opercularis) and the right middle frontal gyrus <u>(Boen et</u> <u>al., 2022; H. Rodrigo et al., 2014)</u>. This pattern suggests that initially students may have relied on previous memories and personal experiences with physical phenomena when attempting to assess plausible trajectories based on the diagrams presented to them. However, at post-instruction, instead of relying on previous personal experiences and contextual information retrieved by the hippocampus, students may have transitioned to using generalized knowledge acquired from physics instruction to assist in suppressing responses for non-Newtonian solutions. While this finding may seem contradictory to our hypotheses regarding the relationship between the hippocampus and constructivist theory, it is important to note that although the hippocampus is critical for initial learning <u>(Schapiro et al., 2019)</u>, there is evidence suggesting that some memory traces become increasingly distributed across the neocortex over time due to repeated activation, transitioning the role of the hippocampus into an index of these cortical representations during memory reactivation <u>(Goode et al., 2020)</u>. However, this may depend on whether detailed or generalized memories are retrieved <u>(Goode et al., 2020)</u>. Consequently, certain abstract and semantic details of memories may become increasingly independent of the hippocampus over time, facilitating representing broader knowledge acquisition, whereas autobiographical and contextually rich memories, as well as certain gist-like memory features may remain hippocampal-dependent <u>(Masís-Obando et al., 2022; Tanaka & McHugh, 2018)</u>. During rest, we observed reduced left intermediate hippocampal connectivity with two clusters in the left precuneus, a region important for episodic memory retrieval and visuospatial processing <u>(Foudil & Macaluso, 2024; Y. Zheng et al., 2023)</u>. Furthermore, our findings showed that the right anterior hippocampus demonstrated reduced functional connectivity with both the left and right lingual gyrus and the left postcentral gyrus at post-instruction. The right hippocampus is typically associated with visuospatial memory, with anterior portions often involved in novelty detection, envisioning future scenarios, and recalling autobiographical visual scenes. Additionally, the lingual gyrus is critical in supporting visual memory, while the left postcentral gyrus has been associated with employing egocentric (self-focused) motor-based strategies during tasks requiring mental rotation <u>(Tomasino & Gremese, 2016)</u>. Combined, decreased connectivity between left intermediate hippocampus and left precuneus, as well as between right anterior hippocampus and bilateral lingual gyrus and left postcentral gyrus, may provide additional evidence of reduced reliance on specific hippocampal functions for certain cognitive processes following formal physics instruction. Initially, for understanding visuospatial physics concepts, students may have relied on previous experiences and intuitive reasoning to facilitate their initial understanding of various physics concepts through the instructional period <u>(Neupärtl et al., 2020)</u>. However, findings from prior research suggest that physics experts tend to focus more on object-object relationships and rely less on metaphors and analogies in their mental representations of problems in their area of expertise in addition to having developed more precise schemata and knowledge structures that allow them to better identify patterns and retrieve relevant information more efficiently <u>(Dixon & Johnson, 2011; Singh, 2008)</u>. Consequently, these resting state findings may indicate a shift from relying on hippocampal-dependent memory processes to facilitate understanding of physics concepts to relying on generalized knowledge distributed across the neocortex that is increasingly independent of the hippocampus.

### Hippocampal Connectivity Changes Preferentially Modulated by Pedagogical Approaches

Results of the interaction analyses indicated that hippocampal functional connectivity may be preferentially preserved and strengthened depending on the pedagogical approach to physics education. During the FCI task, modeling students showed increased connectivity between the right anterior hippocampus and left superior parietal lobule at post-instruction compared to lecture-based students. Additional functions of the right anterior hippocampus, other than spatial information processing, is to support broad, gist-like memory features that facilitate generalization from prior memories, as well as schema-based integration, which incorporates contextual information into newly formed cognitive frameworks <u>(Igarashi et al., 2022;</u> <u>Masís-Obando et al., 2022; Robin & Moscovitch, 2017)</u>. These properties stem from the hippocampus’ role in pattern recognition and extraction of relevant information from partial cues <u>(Igarashi et al., 2022)</u>. Furthermore, the left superior parietal lobule is associated with spatial cognition and visuospatial attention <u>(Wu et al., 2016)</u>. When compared to novices, physics experts are more likely to construct accurate mental representations of problems, visualizing and efficiently extracting relevant physics-related information while attending to important spatial features <u>(Singer, S.R et al., 2012)</u>. Furthermore, as a result of students’ increased engagement with experiential learning and constructivist approaches, modeling instruction may provide multiple sources of information during encoding when learning Newtonian concepts, allowing for more opportunities for partial cue recall <u>(Ross et al., 2018)</u>. When faced with physics problems, important neural circuitry, such as the right anterior hippocampus and left superior parietal lobule, may activate more readily due to partial cue retrieval, helping to identify and apply relevant physics principles dictating the scenarios in the FCI problem. Notably, connectivity between the right anterior hippocampus and left superior parietal lobule did not show substantial correlation with FCI accuracy nor PRI and WMI. Nevertheless, while modeling students may exhibit emerging neural patterns suggestive of increasing expert-like approaches to physics problem solving, introductory physics still may be too early to see tangible benefits of developing problem solving approaches manifest in the FCI task.

For the PK task, modeling students exhibited a greater increase in left anterior hippocampal connectivity with bilateral cerebellum compared to lecture-based students. The PK task primarily centers on retrieval of semantic information and the left anterior hippocampal areas have been demonstrated to increase in task activity related to successful semantic memory <u>(Heckers, 2002)</u>. In addition, both crus 1 and crus 2 have also been associated with semantic processing and language comprehension, as well as the automatic retrieval of words and their associated semantic representations <u>(Nakatani et al., 2022; Petríková et al., 2023)</u>. Since modeling students typically learn in more diverse learning environments, entailing group activities, conversations, as well as engaging in experimental conditions, more context-rich information is available to bind physics terminology with their associated meanings. This interpretation is also further supported by the finding that hippocampal-cerebellar connectivity was associated with increased PK accuracy, emphasizing the role this connection plays in accurate semantic retrieval.

Finally, resting state findings revealed that modeling students showed increased connectivity between left intermediate hippocampus and left superior frontal gyrus, a region important in the mental manipulation and monitoring of information <u>(Boisgueheneuc et al., 2006)</u>. As an active learning approach, modeling instruction is a primarily student-led course, where students are expected to be actively involved in constructing their own understanding of physics concepts, a primary feature of constructivist philosophy. Consequently, students will have periods of monitoring and assessing their own understanding as they collaborate with peers during the learning process to resolve any misconceptions they may have and subsequently update their prior knowledge with more accurate information. Thus, increased connectivity between left intermediate hippocampus and left superior frontal gyrus may reflect that process, consequently highlighting the role of constructivism in active learning approaches. In addition, lecture-based students exhibited increased connectivity between left intermediate hippocampus and right fusiform gyrus, which is associated with facial processing as well as non-verbal associative semantic processing <u>(Mion et al., 2010)</u> at post-instruction. A significant feature of lecture-based learning is observational learning <u>(Bellebaum et al., 2012)</u>, which is the process of being instructed passively and expanding knowledge through observation. Thus, this pattern may be reflective of that process, with conceptual information being learned and understood through observing and listening to their instructor impart information onto them. Interestingly, while lecture-based students exhibited greater left intermediate hippocampus and right fusiform gyrus connectivity, performance on WMI decreased as connectivity between these two regions increased. The fusiform gyrus is involved in working memory <u>(Druzgal & D’Esposito, 2001)</u>; thus, hyperconnectivity between these two regions could disrupt performance by intruding upon cognitive flexibility or rapid information updating, ultimately resulting in poorer task performance <u>(Taghia et al., 2018)</u>.

## Limitations

This study has several limitations that warrant consideration when interpreting its findings. Firstly, our exclusion criteria reduced the initial sample size from 121 to 90 participants. Participants were excluded for various reasons, including failure to return for post-instruction scans, technical difficulties during scanning, and data quality issues flagged by MRIQC (i.e., high mean framewise displacement, high ghost-to-signal ratio, low signal-to-noise ratio, or low entropy focus criterion). Furthermore, our analysis included only participants who completed at least one valid functional run for all three paradigms: FCI task, PK task, and rest. This approach was necessary to ensure that observed differences in hippocampal connectivity primarily reflected longitudinal changes or variations due to instructional methodology, rather than confounded due to comparison of different subgroups across tasks. Importantly, participant information comparisons revealed no significant differences between modeling and lecture-based student groups, either in the original sample or the sub-sample. Secondly, our study employed a static functional connectivity (sFC) analytic approach. While sFC is a common technique for assessing average brain connectivity over an entire scan session, it may obscure dynamic changes. Research findings suggest that connectivity patterns can fluctuate rapidly, potentially on the timescale of seconds. Consequently, sFC methods might mask transient patterns of brain activity relevant to the learning process. Given findings from the main effect of time, it is possible that dynamic functional connectivity analyses may have revealed a brief increase in hippocampal connectivity with various regions at post-instruction that were masked due this FC approach. Consequently, future studies could employ dynamic functional connectivity approaches to investigate how different instructional methods influence time-varying hippocampal connectivity while examining the strength and persistence of these fluctuations throughout the scanning session. Finally, although the study design was longitudinal, it exclusively focused on introductory physics students. Many STEM educational programs often require students to take additional physics courses as part of their core curriculum. Therefore, future research should investigate whether the pedagogy-dependent-induced connectivity patterns observed here intensify in higher-level physics courses as students engage with more complex, standardized physics content.

## Conclusions

Herein, we assessed how different instructional methodologies in an introductory physics course (i.e., active vs. passive learning) was associated with hippocampal whole-brain connectivity at pre- vs. post-instruction through three cognitive contexts: physics perceptual reasoning (i.e., FCI task), physics knowledge retrieval (i.e., PK task), and rest. To this end, we predicted increased hippocampal connectivity at post-instruction regardless of the pedagogical approach. However, we observed decreases in hippocampal connectivity at post-instruction, suggesting that some cognitive processes may across time become increasingly independent of the hippocampus as they become more distributed across the neocortex. Moreover, we also predicted that active learning students would exhibit more robust increased hippocampal connectivity changes from pre- to post-instruction. In terms of pedagogical differences, active learning students exhibited stronger hippocampal connectivity with regions associated with visuospatial processing, semantic processing, and self-monitoring, highlighting the role of constructivist approaches in active learning. Lecture-based students exhibited increased hippocampal-fusiform connectivity at post-instruction, reflecting their emphasis on understanding conceptual knowledge through passive observational approaches. Overall, these findings suggest that while some cognitive processes may become increasingly hippocampal-independent with physics learning, others become more hippocampal-dependent, although this depends on the pedagogical approach. Ultimately, the findings of this study suggest that constructivism is central to active learning teaching methodologies, which may be reflected in the observed changes in hippocampal connectivity.

## Acknowledgments

Data collection for this project was funded by NSF REAL DRL-1420627 (ARL, EB, SMP, JEB). Contributions from co-authors were provided with support from NIH U01-DA041156 (ARL, MTS, MCR, KLB). Additional support was provided by the Office of Research and Economic Development and the Dissertation Fellowship from the University Graduate School at Florida International University (FIU). Special thanks to the FIU Instructional & Research Computing Center (IRCC, http://ircc.fiu.edu) for providing the HPC and computing resources that contributed to the research results reported within this paper, as well as to the Department of Psychology of the University of Miami for providing access to their MRI scanner. Additional thanks to Dr. Jeremy Elman for sharing semantic retrieval task stimuli. Lastly, the authors would like to thank the FIU undergraduate students who volunteered, participated, and contributed to this project.

## Data Availability

All tabular (behavioral) data and the statistical brain volumes used to perform these analyses (i.e., Z-score contrast maps from the main effect of time, main effect of class, and interaction effects) are available on the Open Science Framework (OSF) project page at https://osf.io/myv5x/.

## Code Availability

The <u>Plachti et al., 2019</u> parcellation is available from via the ANIMA database at https://anima.fz-juelich.de/studies/Plachti_Hippocampus_2019. Scripts from the statistical analyses and figures used in this study are available on the OSF project page at https://osf.io/myv5x/.

## Author Contributions

DDS and ARL conceived and designed the study. JEB acquired data. DDS analyzed the data. DDS, JAP, KLB, MCR, and TS contributed scripts, pipelines, and figures. DDS and ARL wrote the paper, and all authors contributed to the revisions and approved the final version.

## References

Anand, K., & Dhikav, V. (2012). Hippocampus in health and disease: An overview. Annals of Indian Academy of Neurology, 15(4), 239. 10.4103/0972-2327.104323

Atkinson, D., Hill, D. L. G., Stoyle, P. N. R., Summers, P. E., & Keevil, S. F. (1997). Automatic correction of motion artifacts in magnetic resonance images using an entropy focus criterion. IEEE Transactions on Medical Imaging, 16(6), 903–910. 10.1109/42.650886

Avants, B., Epstein, C., Grossman, M., & Gee, J. (2008). Symmetric diffeomorphic image registration with cross-correlation: Evaluating automated labeling of elderly and neurodegenerative brain. Medical Image Analysis, 12(1), 26–41. 10.1016/j.media.2007.06.004

Bartley, J. E., Riedel, M. C., Salo, T., Boeving, E. R., Bottenhorn, K. L., Bravo, E. I., Odean, R., Nazareth, A., Laird, R. W., Sutherland, M. T., Pruden, S. M., Brewe, E., & Laird, A. R. (2019). Brain activity links performance in science reasoning with conceptual approach. Npj Science of Learning, 4(1), 20. 10.1038/s41539-019-0059-8

Behzadi, Y., Restom, K., Liau, J., & Liu, T. T. (2007). A component based noise correction method (CompCor) for BOLD and perfusion based fMRI. NeuroImage, 37(1), 90–101. 10.1016/j.neuroimage.2007.04.042

Bein, O., Reggev, N., & Maril, A. (2020). Prior knowledge promotes hippocampal separation but cortical assimilation in the left inferior frontal gyrus. Nature Communications, 11(1), 4590. 10.1038/s41467-020-18364-1

Bellebaum, C., Jokisch, D., Gizewski, E. R., Forsting, M., & Daum, I. (2012). The neural coding of expected and unexpected monetary performance outcomes: Dissociations between active and observational learning. Behavioural Brain Research, 227(1), 241–251. 10.1016/j.bbr.2011.10.042

Blanquat, P. D. S., Hok, V., Save, E., Poucet, B., & Chaillan, F. A. (2013). Differential role of the dorsal hippocampus, ventro-intermediate hippocampus, and medial prefrontal cortex in updating the value of a spatial goal. Hippocampus, 23(5), 342–351. 10.1002/hipo.22094

Boisgueheneuc, F. D., Levy, R., Volle, E., Seassau, M., Duffau, H., Kinkingnehun, S., Samson, Y., Zhang, S., & Dubois, B. (2006). Functions of the left superior frontal gyrus in humans: A lesion study. Brain, 129(12), 3315–3328. 10.1093/brain/awl244

Borders, A. A., Aly, M., Parks, C. M., & Yonelinas, A. P. (2017). The hippocampus is particularly important for building associations across stimulus domains. Neuropsychologia, 99, 335–342. 10.1016/j.neuropsychologia.2017.03.032

Brewe, E., Bartley, J. E., Riedel, M. C., Sawtelle, V., Salo, T., Boeving, E. R., Bravo, E. I., Odean, R., Nazareth, A., Bottenhorn, K. L., Laird, R. W., Sutherland, M. T., Pruden, S. M., & Laird, A. R. (2018). Toward a Neurobiological Basis for Understanding Learning in University Modeling Instruction Physics Courses. Frontiers in ICT, 5, 10. 10.3389/fict.2018.00010

Burgess, N., Maguire, E. A., & O’Keefe, J. (2002). The Human Hippocampus and Spatial and Episodic Memory. Neuron, 35(4), 625–641. 10.1016/S0896-6273(02)00830-9

Buxton, R. B., Uludağ, K., Dubowitz, D. J., & Liu, T. T. (2004). Modeling the hemodynamic response to brain activation. NeuroImage, 23, S220–S233. 10.1016/j.neuroimage.2004.07.013

Chu, H.-E., Treagust, D. F., & Chandrasegaran, A. L. (2008). Naïve Students’ Conceptual Development and Beliefs: The Need for Multiple Analyses to Determine what Contributes to Student Success in a University Introductory Physics Course. Research in Science Education, 38(1), 111–125. 10.1007/s11165-007-9068-3

Cole, M. W., Ito, T., Bassett, D. S., & Schultz, D. H. (2016). Activity flow over resting-state networks shapes cognitive task activations. Nature Neuroscience, 19(12), 1718–1726. 10.1038/nn.4406

Dale, A. M., Fischl, B., & Sereno, M. I. (1999). Cortical Surface-Based Analysis. NeuroImage, 9(2), 179–194. 10.1006/nimg.1998.0395

Dancy, M., Henderson, C., Apkarian, N., Johnson, E., Stains, M., Raker, J. R., & Lau, A. (2024). Physics instructors’ knowledge and use of active learning has increased over the last decade but most still lecture too much. Physical Review Physics Education Research, 20(1), 010119. 10.1103/PhysRevPhysEducRes.20.010119

Das, A., & Menon, V. (2021). Asymmetric Frequency-Specific Feedforward and Feedback Information Flow between Hippocampus and Prefrontal Cortex during Verbal Memory Encoding and Recall. The Journal of Neuroscience, 41(40), 8427–8440. 10.1523/JNEUROSCI.0802-21.2021

Dixon, R. A., & Johnson, S. D. (2011). Experts vs. Novices: Differences in How Mental Representations are Used in Engineering Design. Journal of Technology Education, 23(1). 10.21061/jte.v23i1.a.5

Druzgal, T. J., & D’Esposito, M. (2001). Activity in fusiform face area modulated as a function of working memory load. Cognitive Brain Research, 10(3), 355–364. 10.1016/S0926-6410(00)00056-2

Elman, J. A., Klostermann, E. C., Marian, D. E., Verstaen, A., & Shimamura, A. P. (2012). Neural correlates of metacognitive monitoring during episodic and semantic retrieval. *Cognitive, Affective*, & Behavioral Neuroscience, 12(3), 599–609. 10.3758/s13415-012-0096-8

Esteban, O., Birman, D., Schaer, M., Koyejo, O. O., Poldrack, R. A., & Gorgolewski, K. J. (2017). MRIQC: Advancing the automatic prediction of image quality in MRI from unseen sites. PLOS ONE, 12(9), e0184661. 10.1371/journal.pone.0184661

Esteban, O., Ciric, R., Finc, K., Blair, R. W., Markiewicz, C. J., Moodie, C. A., Kent, J. D., Goncalves, M., DuPre, E., Gomez, D. E. P., Ye, Z., Salo, T., Valabregue, R., Amlien, I. K., Liem, F., Jacoby, N., Stojić, H., Cieslak, M., Urchs, S., … Gorgolewski, K. J. (2020). Analysis of task-based functional MRI data preprocessed with fMRIPrep. Nature Protocols, 15(7), 2186–2202. 10.1038/s41596-020-0327-3

Fanselow, M. S., & Dong, H.-W. (2010). Are the Dorsal and Ventral Hippocampus Functionally Distinct Structures? Neuron, 65(1), 7–19. 10.1016/j.neuron.2009.11.031

Foudil, S.-A., & Macaluso, E. (2024). The influence of the precuneus on the medial temporal cortex determines the subjective quality of memory during the retrieval of naturalistic episodes. Scientific Reports, 14(1), 7943. 10.1038/s41598-024-58298-y

Freeman, S., Eddy, S. L., McDonough, M., Smith, M. K., Okoroafor, N., Jordt, H., & Wenderoth, M. P. (2014). Active learning increases student performance in science, engineering, and mathematics. Proceedings of the National Academy of Sciences, 111(23), 8410–8415. 10.1073/pnas.1319030111

Fuchsberger, T., & Paulsen, O. (2022). Modulation of hippocampal plasticity in learning and memory. Current Opinion in Neurobiology, 75, 102558. 10.1016/j.conb.2022.102558

Goode, T. D., Tanaka, K. Z., Sahay, A., & McHugh, T. J. (2020). An Integrated Index: Engrams, Place Cells, and Hippocampal Memory. Neuron, 107(5), 805–820. 10.1016/j.neuron.2020.07.011

Gorgolewski, K., Burns, C. D., Madison, C., Clark, D., Halchenko, Y. O., Waskom, M. L., & Ghosh, S. S. (2011). Nipype: A Flexible, Lightweight and Extensible Neuroimaging Data Processing Framework in Python. Frontiers in Neuroinformatics, 5. 10.3389/fninf.2011.00013

Greve, D. N., & Fischl, B. (2009). Accurate and robust brain image alignment using boundary-based registration. NeuroImage, 48(1), 63–72. 10.1016/j.neuroimage.2009.06.060

Hartman, D. E. (2009). Wechsler Adult Intelligence Scale IV (WAIS IV): Return of the Gold Standard. Applied Neuropsychology, 16(1), 85–87. 10.1080/09084280802644466

Heckers, S. (2002). Hippocampal and Brain Stem Activation during Word Retrieval after Repeated and Semantic Encoding. Cerebral Cortex, 12(9), 900–907. 10.1093/cercor/12.9.900

Hefter, M. H., Fromme, B., & Berthold, K. (2022). Digital training intervention on strategies for tackling physical misconceptions—Self-explanation matters. Applied Cognitive Psychology, 36(3), 648–658. 10.1002/acp.3951

Hestenes, D., Wells, M., & Swackhamer, G. (1992). Force concept inventory. The Physics Teacher, 30(3), 141–158. 10.1119/1.2343497

Igarashi, K. M., Lee, J. Y., & Jun, H. (2022). Reconciling neuronal representations of schema, abstract task structure, and categorization under cognitive maps in the entorhinal-hippocampal-frontal circuits. Current Opinion in Neurobiology, 77, 102641. 10.1016/j.conb.2022.102641

Johari, A. H. & Muslim. (2018). Application of experiential learning model using simple physical kit to increase attitude toward physics student senior high school in fluid. Journal of Physics: Conference Series, 1013, 012032. 10.1088/1742-6596/1013/1/012032

Kafkas, A., & Montaldi, D. (2018). How do memory systems detect and respond to novelty? Neuroscience Letters, 680, 60–68. 10.1016/j.neulet.2018.01.053

Kotsis, K. T. (2024). Correcting Students’ Misconceptions in Physics Using Experiments Designed by ChatGPT. European Journal of Contemporary Education and E-Learning, 2(2), 83–100. 10.59324/ejceel.2024.2(2).07

Lasry, N., Rosenfield, S., Dedic, H., Dahan, A., & Reshef, O. (2011). The puzzling reliability of the Force Concept Inventory. American Journal of Physics, 79(9), 909–912. 10.1119/1.3602073

Luo, X., Rechardt, A., Sun, G., Nejad, K. K., Yáñez, F., Yilmaz, B., Lee, K., Cohen, A. O., Borghesani, V., Pashkov, A., Marinazzo, D., Nicholas, J., Salatiello, A., Sucholutsky, I., Minervini, P., Razavi, S., Rocca, R., Yusifov, E., Okalova, T., … Love, B. C. (2024). Large language models surpass human experts in predicting neuroscience results. Nature Human Behaviour, 9(2), 305–315. 10.1038/s41562-024-02046-9

Maguire, E. A. (2014). Memory consolidation in humans: New evidence and opportunities. Experimental Physiology, 99(3), 471–486. 10.1113/expphysiol.2013.072157

Masís-Obando, R., Norman, K. A., & Baldassano, C. (2022). Schema representations in distinct brain networks support narrative memory during encoding and retrieval. eLife, 11, e70445. 10.7554/eLife.70445

Md Rashid, S. N., Makhtar, N., Jaafar, N. F., & Abdullah, S. Z. (2024). Improving Students’ Understanding in Physics Using Experiential Learning. International Journal of Academic Research in Progressive Education and Development, 13(1), Pages 1256–1262. 10.6007/IJARPED/v13-i1/20625

Mion, M., Patterson, K., Acosta-Cabronero, J., Pengas, G., Izquierdo-Garcia, D., Hong, Y. T., Fryer, T. D., Williams, G. B., Hodges, J. R., & Nestor, P. J. (2010). What the left and right anterior fusiform gyri tell us about semantic memory. Brain, 133(11), 3256–3268. 10.1093/brain/awq272

Mughal, F., & Zafar, A. (2011). Experiential Learning from a Constructivist Perspective: Reconceptualizing the Kolbian Cycle. International Journal of Learning and Development, 1(2), 27. 10.5296/ijld.v1i2.1179

Nakatani, H., Nakamura, Y., & Okanoya, K. (2022). Respective Involvement of the Right Cerebellar Crus I and II in Syntactic and Semantic Processing for Comprehension of Language. The Cerebellum, 22(4), 739–755. 10.1007/s12311-022-01451-y

Neupärtl, N., Tatai, F., & Rothkopf, C. A. (2020). Intuitive physical reasoning about objects’ masses transfers to a visuomotor decision task consistent with Newtonian physics. PLOS Computational Biology, 16(10), e1007730. 10.1371/journal.pcbi.1007730

Pande, M., & Bharathi, S. V. (2020). Theoretical foundations of design thinking – A constructivism learning approach to design thinking. Thinking Skills and Creativity, 36, 100637. 10.1016/j.tsc.2020.100637

Peraza, J. A., Kent, J. D., Nichols, T. E., Poline, J.-B., De La Vega, A., & Laird, A. R. (2025). NiCLIP: Neuroimaging contrastive language-image pretraining model for predicting text from brain activation images. 10.1101/2025.06.14.659706

Petríková, D., Marko, M., Rovný, R., & Riečanský, I. (2023). Electrical stimulation of the cerebellum facilitates automatic but not controlled word retrieval. Brain Structure and Function, 228(9), 2137–2146. 10.1007/s00429-023-02712-0

Plachti, A., Eickhoff, S. B., Hoffstaedter, F., Patil, K. R., Laird, A. R., Fox, P. T., Amunts, K., & Genon, S. (2019). Multimodal Parcellations and Extensive Behavioral Profiling Tackling the Hippocampus Gradient. Cerebral Cortex, 29(11), 4595–4612. 10.1093/cercor/bhy336

Poldrack, R. A., Kittur, A., Kalar, D., Miller, E., Seppa, C., Gil, Y., Parker, D. S., Sabb, F. W., & Bilder, R. M. (2011). The Cognitive Atlas: Toward a Knowledge Foundation for Cognitive Neuroscience. Frontiers in Neuroinformatics, 5. 10.3389/fninf.2011.00017

Porter, A. M., Chu, R. Y., & Ivie, R. (2024). *Attrition and Persistence in Undergraduate Physics Programs* (0 ed.). American Institute of Physics. 10.1063/sr.a213485edb

Reuter, M., Rosas, H. D., & Fischl, B. (2010). Highly accurate inverse consistent registration: A robust approach. NeuroImage, 53(4), 1181–1196. 10.1016/j.neuroimage.2010.07.020

Robin, J., & Moscovitch, M. (2017). Details, gist and schema: Hippocampal–neocortical interactions underlying recent and remote episodic and spatial memory. Current Opinion in Behavioral Sciences, 17, 114–123. 10.1016/j.cobeha.2017.07.016

Robinson, J. L., Salibi, N., & Deshpande, G. (2016). Functional connectivity of the left and right hippocampi: Evidence for functional lateralization along the long-axis using meta-analytic approaches and ultra-high field functional neuroimaging. NeuroImage, 135, 64–78. 10.1016/j.neuroimage.2016.04.022

Roche, J., Self, B., & Widmann, J. (2017). Student Self-Explanation When Solving a Rigid Body Kinetics Concept Question. 2017 Pacific Southwest Section Meeting Proceedings, 29235. 10.18260/1-2--29235

Ross, D. A., Sadil, P., Wilson, D. M., & Cowell, R. A. (2018). Hippocampal Engagement during Recall Depends on Memory Content. Cerebral Cortex, 28(8), 2685–2698. 10.1093/cercor/bhx147

Sandrone, S., Scott, G., Anderson, W. J., & Musunuru, K. (2021). Active learning-based STEM education for in-person and online learning. Cell, 184(6), 1409–1414. 10.1016/j.cell.2021.01.045

Satterthwaite, T. D., Elliott, M. A., Gerraty, R. T., Ruparel, K., Loughead, J., Calkins, M. E., Eickhoff, S. B., Hakonarson, H., Gur, R. C., Gur, R. E., & Wolf, D. H. (2013). An improved framework for confound regression and filtering for control of motion artifact in the preprocessing of resting-state functional connectivity data. NeuroImage, 64, 240–256. 10.1016/j.neuroimage.2012.08.052

Schapiro, A. C., Reid, A. G., Morgan, A., Manoach, D. S., Verfaellie, M., & Stickgold, R. (2019). The hippocampus is necessary for the consolidation of a task that does not require the hippocampus for initial learning. Hippocampus, 29(11), 1091–1100. 10.1002/hipo.23101

Shafer, D., Girotti-Hernandez, D., & Stelzer, T. (2024). Evolving study strategies and support structures of introductory physics students. Physical Review Physics Education Research, 20(2), 020114. 10.1103/PhysRevPhysEducRes.20.020114

Singer, S.R, Nielsen, N., & Schweingruber, H. A. (2012). Discipline-Based Education Research: Understanding and Improving Learning in Undergraduate Science and Engineering (p. 13362). National Academies Press. 10.17226/13362

Singh, C. (2008). Assessing student expertise in introductory physics with isomorphic problems. I. Performance on nonintuitive problem pair from introductory physics. Physical Review Special Topics - Physics Education Research, 4(1), 010104. 10.1103/PhysRevSTPER.4.010104

Smith, D. D., Meca, A., Bottenhorn, K. L., Bartley, J. E., Riedel, M. C., Salo, T., Peraza, J. A., Laird, R. W., Pruden, S. M., Sutherland, M. T., Brewe, E., & Laird, A. R. (2023). Task-based attentional and default mode connectivity associated with science and math anxiety profiles among university physics students. Trends in Neuroscience and Education, 32, 100204. 10.1016/j.tine.2023.100204

Sokoloff, D. R., Laws, P. W., & Thornton, R. K. (2007). *RealTime Physics*: Active learning labs transforming the introductory laboratory. European Journal of Physics, 28(3), S83–S94. 10.1088/0143-0807/28/3/S08

Strange, B. A., Witter, M. P., Lein, E. S., & Moser, E. I. (2014). Functional organization of the hippocampal longitudinal axis. Nature Reviews Neuroscience, 15(10), 655–669. 10.1038/nrn3785

Taghia, J., Cai, W., Ryali, S., Kochalka, J., Nicholas, J., Chen, T., & Menon, V. (2018). Uncovering hidden brain state dynamics that regulate performance and decision-making during cognition. Nature Communications, 9(1), 2505. 10.1038/s41467-018-04723-6

Tanaka, K. Z., & McHugh, T. J. (2018). The Hippocampal Engram as a Memory Index. Journal of Experimental Neuroscience, 12, 1179069518815942. 10.1177/1179069518815942

Tomasino, B., & Gremese, M. (2016). Effects of Stimulus Type and Strategy on Mental Rotation Network: An Activation Likelihood Estimation Meta-Analysis. Frontiers in Human Neuroscience, 9. 10.3389/fnhum.2015.00693

Tustison, N. J., Avants, B. B., Cook, P. A., Yuanjie Zheng, Egan, A., Yushkevich, P. A., & Gee, J. C. (2010). N4ITK: Improved N3 Bias Correction. IEEE Transactions on Medical Imaging, 29(6), 1310–1320. 10.1109/TMI.2010.2046908

Von Korff, J., Archibeque, B., Gomez, K. A., Heckendorf, T., McKagan, S. B., Sayre, E. C., Schenk, E. W., Shepherd, C., & Sorell, L. (2016). Secondary analysis of teaching methods in introductory physics: A 50 k-student study. American Journal of Physics, 84(12), 969–974. 10.1119/1.4964354

Wang, L., Zhang, J., Liu, T., Chen, D., Yang, D., Go, R., Wu, J., & Yan, T. (2022). Prediction of Cognitive Task Activations via Resting-State Functional Connectivity Networks: An EEG Study. IEEE Transactions on Cognitive and Developmental Systems, 14(1), 181–188. 10.1109/TCDS.2020.3031604

Wechsler, D. (2012). Wechsler Adult Intelligence Scale—Fourth Edition [Dataset]. American Psychological Association (APA). 10.1037/t15169-000

Wiltgen, B. J., Zhou, M., Cai, Y., Balaji, J., Karlsson, M. G., Parivash, S. N., Li, W., & Silva, A. J. (2010). The Hippocampus Plays a Selective Role in the Retrieval of Detailed Contextual Memories. Current Biology, 20(15), 1336–1344. 10.1016/j.cub.2010.06.068

Wittkuhn, L., & Schuck, N. W. (2021). Dynamics of fMRI patterns reflect sub-second activation sequences and reveal replay in human visual cortex. Nature Communications, 12(1), 1795. 10.1038/s41467-021-21970-2

Wu, Y., Wang, J., Zhang, Y., Zheng, D., Zhang, J., Rong, M., Wu, H., Wang, Y., Zhou, K., & Jiang, T. (2016). The Neuroanatomical Basis for Posterior Superior Parietal Lobule Control Lateralization of Visuospatial Attention. Frontiers in Neuroanatomy, 10. 10.3389/fnana.2016.00032

Zeidman, P., & Maguire, E. A. (2016). Anterior hippocampus: The anatomy of perception, imagination and episodic memory. Nature Reviews Neuroscience, 17(3), 173–182. 10.1038/nrn.2015.24

Zhang, Y., Brady, M., & Smith, S. (2001). Segmentation of brain MR images through a hidden Markov random field model and the expectation-maximization algorithm. IEEE Transactions on Medical Imaging, 20(1), 45–57. 10.1109/42.906424

Zheng, A., Montez, D. F., Marek, S., Gilmore, A. W., Newbold, D. J., Laumann, T. O., Kay, B. P., Seider, N. A., Van, A. N., Hampton, J. M., Alexopoulos, D., Schlaggar, B. L., Sylvester, C. M., Greene, D. J., Shimony, J. S., Nelson, S. M., Wig, G. S., Gratton, C., McDermott, K. B., … Dosenbach, N. U. F. (2021). Parallel hippocampal-parietal circuits for self- and goal-oriented processing. Proceedings of the National Academy of Sciences, 118(34), e2101743118. 10.1073/pnas.2101743118

Zheng, Y., Xie, L., Huang, Z., Peng, J., Huang, S., Guo, R., Huang, J., Lin, Z., Zhuang, Z., Yin, J., Hou, Z., & Ma, S. (2023). Enhanced activity of the left precuneus as a predictor of visuospatial dysfunction correlates with disease activity in rheumatoid arthritis. European Journal of Medical Research, 28(1), 276. 10.1186/s40001-023-01224-1

