## Supplemental Information for "Hippocampal functional connectivity changes associated with active and lecture-based physics learning"

This document includes:

- Supplemental Methods
- Supplemental Results
- Supplemental Table 1
- Supplemental Table 2
- Supplemental Table 3
- Supplemental References

### SUPPLEMENTAL METHODS

#### Neuroimaging Preprocessing

All functional magnetic resonance imaging (fMRI) data used in this study were preprocessed using fMRIPrep 1.5.0, a BIDS-compliant Python package which automatically provides a high-quality, reproducible preprocessing workflow with minimal user intervention (Esteban et al., 2019).

##### Anatomical Data Preprocessing

T1-weighted images were corrected for intensity non-uniformity (INU) with N4FieldCorrection (Tustison et al., 2010), distributed with ANTs 2.2.0 (Avants et al., 2008, RRID:SCR_004757). The T1w-reference was then skull-stripped with a Nipype implementation of the antsBrainExtraction.sh workflow (from ANTs), using OASIS30ANTs as target template. Brain tissue segmentation of the cerebrospinal fluid (CSF), white-matter (WM) and gray-matter (GM) was performed on the brain-extracted T1w using fast (FSL 5.0.9, RRIDLSCR_002823, Zhang et al., 2001). A T1w-reference map was computed after registration of 2 T1w images (after INU-correction) using mri_robust_template (FreeSurfer 6.0.1, Reuter et al., 2010). Brain surfaces were reconstructed using recon-all (FreeSurfer 6.0.1, RRID:SCR_001847, Dale et al., 1999), and the brain mask estimated previously was refined with a custom variation of the method to reconcile ANTs-derived and FreeSurfer-derived segmentations of the cortical gray-matter of Mindboggle (RRID:SCR_002438, Klein et al., 2017). Volume-based spatial normalization to one standard space (MNI152NLin2009cAsym) was performed through nonlinear registration with antsRegistration (ANTs 2.2.0), using brain-extracted version of both T1w reference and the T1w template. The following template was selected for spatial normalization: ICBM 152 Nonlinear Asymmetrical template version 2009c {Fonov et al. (2009), RRID:SCR_008796: TemplateFlow ID: MNI152NLin2009cAsym].

##### Functional Data Preprocessing

Each participant's dataset contained 1-3 runs of task-based functional magnetic resonance imaging (fMRI) data. For each BOLD run found per subject (across all tasks and sessions), the following preprocessing was performed. First, a reference volume and its skull-stripped version were generated using a custom methodology of fMRIPrep. A deformation field to correct for susceptibility distortions was estimated based on fMRIPrep's fieldmap-*less* approach. The deformation field is that resulting from co-registering the BOLD reference to the same-subject T1w-reference with its intensity inverted (Huntenburg, 2014; Wang et al., 2017). Registration is performed with antsRegistration (ANTs 2.2.0), and the process regularized by constraining deformation to be nonzero only along the phase-encoding direction, and modulated with an average fieldmap template (Treiber et al., 2016). Based on the estimated susceptibility distortion, an unwarped BOLD reference was calculated for a more accurate co-registration with the anatomical reference. The BOLD reference was then co-registered to the T1w reference using bbregister (FreeSurfer) which implements boundary-based registration (Greve & Fischl, 2009). Co-registration was configured with six degrees of freedom. Head-motion parameters with respect to the BOLD reference (transformation matrices, and six corresponding rotation and translation parameters) are estimated before any spatiotemporal filtering using mcflirt (FSL 5.0.9, Jenkinson et al., 2002). The BOLD time-series were resampled to surfaces on the following spaces: *fsaverage5*. The BOLD time-series (including slice-timing correction when applied) were resampled onto their original, native space by applying a single, composite transform to correct for head-motion and susceptibility distortions. These resampled BOLD time-series will be referred to as preprocessed BOLD in original space, or just preprocessed BOLD. The BOLD time-series were resampled into standard space, generating a preprocessed BOLD run in *MNI152NLin2009cAsym* space. First, a reference volume and its skull-stripped version were generated using a custom methodology of fMRIPrep. Several confounding time-series were calculated based on the preprocessed BOLD: framewise displacement (FD), DVARS and three region-wise global signals. FD and DVARS are calculated for each functional run, both using their implementations in `Nipype` (following the definitions by Power et al. 2014). The three global signals are extracted within the CSF, the WM, and the whole-brain masks. Additionally, a set of physiological regressors were extracted to allow for component-based noise correction (CompCor, Behzadi et al. 2007). Principal components are estimated after high-pass filtering the preprocessed BOLD time-series (using a discrete cosine filter with 128s cut-off) for the two CompCor variants: temporal (tCompCor) and anatomical (aCompCor). tCompCor components are then calculated from the top 5% variable voxels within a mask covering the subcortical regions. This subcortical mask is obtained by heavily eroding the brain mask, which ensures it does not include cortical GM regions. For aCompCor, components are calculated within the intersection of the aforementioned mask and the union of CSF and WM masks calculated in T1w space, after their projection to the native space of each functional run (using the inverse BOLD-to-T1w transformation). Components are also calculated separately within the WM and CSF masks. For each CompCor decomposition, the *k* components with the largest singular values are retained, such that the retained components' time series are sufficient to explain 50 percent of variance across the nuisance mask (CSF, WM, combined, or temporal). The remaining components are dropped from consideration. The head-motion estimates calculated in the correction step were also placed within the corresponding confounds file. The confound time series derived from head motion estimates and global signals were expanded with the inclusion of temporal derivatives and quadratic terms for each (Satterthwaite et al., 2013). Frames that exceeded a threshold of 0.5 mm FD or 1.5 standardised DVARS were annotated as motion outliers. All resamplings can be performed with a single interpolation step by composing all the pertinent transformations (i.e. head-motion transform matrices, susceptibility distortion correction when available, and co-registrations to anatomical and output spaces). Gridded (volumetric) resamplings were performed using `antsApplyTransforms` (`ANTs`), configured with Lanczos interpolation to minimize the smoothing effects of other kernels (Lanczos, 1964). Non-gridded (surface) resamplings were performed using `mri_vol2surf` (`FreesSurfer`).

#### SUPPLEMENTAL RESULTS

Prior to the CAPs analysis, functional images were excluded from the time series extraction process if MRIQC reports indicated that the mean framewise displacement, ghost-to-signal ratio, or entropy focus criterion exceeded the 99th percentile, or if the signal-to-noise ratio was below the 1st percentile; thus, indicating that these runs were outliers. Participants were also excluded if they did not possess at least one functional run for both pre-instruction and post-instruction resting state and task (FCI & PK) data. Based on these criteria, the sample size was reduced from 121 to 91 participants. Consequently, **Supplemental Table 1** presents the demographic information for the active learning (modeling-instruction) and lecture-based instruction groups after applying these exclusion criteria. As stated in the main manuscript, no significant differences between any of the demographic variables were noted between the active learning class or the lecture-based instruction class.

*Table S1.* **Participant Demographic Information After Exclusion Criteria.**

| | Modeling Instruction (*n*=46) | | Lecture-Based Instruction (*n*=44) | |
|---|---|---|---|---|
| | N | Percentage | N | Percentage |
| **Gender** | | | | |
| Male | 25 | 46 | 23 | 48 |
| Female | 21 | 54 | 21 | 52 |
| **Ethnicity** | | | | |
| Hispanic | 34 | 74 | 31 | 70 |
| Non-Hispanic | 12 | 26 | 13 | 30 |
| **Household Income** | | | | |
| < $15,000 | 8 | 17 | 9 | 20 |
| $15,000 - $34,999 | 10 | 22 | 10 | 23 |
| $35,000 - $49,999 | 5 | 11 | 6 | 14 |
| $50,000 - $74,999 | 12 | 26 | 5 | 11 |
| $75,000 - $99,999 | 9 | 20 | 6 | 14 |
| >$100,000 | 2 | 4 | 8 | 18 |
| **Years Enrolled** | | | | |
| Freshman | 3 | 6 | 5 | 11 |
| Sophomore | 24 | 52 | 20 | 45 |
| Junior | 14 | 20 | 12 | 27 |
| Senior | 5 | 11 | 7 | 16 |
| | **Grand Mean** | **Std. Dev.** | **Grand Mean** | **Std. Dev.** |
| **Age** | 19.96 | 1.76 | 19.89 | 1.50 |
| **GPA** | 3.31 | 0.62 | 3.25 | 0.41 |

*Note.* The "N" column represents the sample size of the group, and the "Percentage" column represents the percentage of participants in that group for each categorical variable (i.e., Gender, Ethnicity, Household Income, and Years Enrolled).

We conducted a comprehensive analysis of the demographic characteristics of students enrolled in the active learning classrooms (n=61) versus those in the lecture-based classrooms (n=60). The variables examined included age, sex, ethnicity, household income, grade point average (GPA), and academic standing at FIU (freshman, sophomore, junior, or senior). As detailed in **Supplementary Table 2**, statistical tests revealed no significant differences between the two groups across these demographics. Additionally, we focused on a subset of participants who completed at least one functional run for all tasks and sessions. This subsample comprised 46 students from the active learning classrooms and 44 from the lecture-based classrooms. Consistent with the broader sample, no significant demographic differences were observed between these groups (**Supplementary Table 2**)

**Table S2. Statistics for Demographic Information.**

| | Full Sample (n=121) | | Subsample (n=90) | |
|---|---|---|---|---|
| | Statistic | P-value | Statistic | P-value |
| **Sex** | $X^2(1) = 1.389$ | 0.239 | $X^2(1) = 0.174$ | 0.677 |
| **Ethnicity** | $X^2(1) = 0.00$ | 1.00 | $X^2(1) = 0.017$ | 0.896 |
| **Income** | $X^2(5) = 6.646$ | 0.248 | $X^2(5) = 7.191$ | 0.207 |
| **Years Enrolled** | $X^2(3) = 0.294$ | 0.961 | $X^2(3) = 1.307$ | 0.728 |
| **Pre-Instruction Age** | $t(116.84) = 0.362$ | 0.719 | $t(86.194) = 0.213$ | 0.832 |
| **Post-Instruction Age** | $t(117.73) = 0.301$ | 0.764 | $t(87.223) = 0.254$ | 0.800 |
| **Pre-Instruction GPA** | $t(95.319) = -0.068$ | 0.945 | $t(84.744) = 0.851$ | 0.397 |
| **Post-Instruction GPA** | $t(97.506) = 0.086$ | 0.932 | $t(73.558) = 0.266$ | 0.791 |

After identifying significant clusters that survived correction, we extracted the beta coefficients from the statistical maps, which represent the strength in connectivity between a hippocampal node and brain region. These beta coefficients were then used in subsequent behavioral analyses to assess the correlation between changes in hippocampal-region connectivity and changes in WAIS-IV scores for specific indices (PRI and WMI for the FCI task, VCI and WMI for the PK task, and all indices for resting state) and accuracy on the FCI and PK tasks. Furthermore, for clusters that showed significant contrasts for the main effect of time contrast, the tested beta coefficient was the difference in the beta coefficient (post-instruction - pre-instruction). This allowed us to assess how changes in pre-instruction to post-instruction connectivity were associated with changes in pre-instruction to post-instruction WAIS-IV scores. Additionally, for the interaction contrast, the difference in beta coefficient across time was interacted with instructional methodology (with the modeling instruction group used as the reference). This allowed us to assess how changes in pre-instruction to post-instruction connectivity, moderated by the instructional methodology, were associated with changes in scores from pre-instruction to post-instruction WAIS-IV. As demonstrated in **Supplementary Table 3**, the behavioral analyses revealed only significant interaction results, with the modeling instruction class showing increased PK accuracy associated with greater connectivity between the left anterior hippocampus and the left cerebellum (crus 2) during the PK task. Additionally, this group exhibited enhanced performance on the WAIS-IV WMI correlated with increasing resting state connectivity between the left intermediate hippocampus and the right fusiform gyrus.

**Table S2. Statistics for Behavioral Analysis.**

| Contrast | Hippocampal Seed | Region Label | Term Tested (β Units) | Behavioral Variable | Model Beta Coefficient (β) | P-value |
|---|---|---|---|---|---|---|
| **FCI Task** | | | | | | |
| Main Effect of Time | L. Intermediate Hippocampus | Right Inferior Frontal Gyrus (p. Opercularis) | Δ Connectivity | ΔWAIS-IV PRI | -0.163 | 0.877 |
| | | | | ΔWAIS-IV WMI | 0.735 | 0.646 |
| | | | | ΔMean FCI Accuracy | -0.031 | 0.232 |
| | | Right Middle Frontal Gyrus | | ΔWAIS-IV PRI | -0.394 | 0.627 |

| | | | | | | |
|---|---|---|---|---|---|---|
| | | | | ΔWAIS-IV WMI | 0.616 | 0.617 |
| | | | | ΔMean FCI Accuracy | -0.020 | 0.314 |
| Interaction | R. Anterior Hippocampus | Left Superior Parietal Lobule | Δ Connectivity x Instructional Methodology | ΔWAIS-IV PRI | 0.458 | 0.838 |
| | | | | ΔWAIS-IV WMI | -4.362 | 0.197 |
| | | | | ΔMean FCI Accuracy | 0.087 | 0.110 |
| **PK Task** | | | | | | |
| Interaction | L. Anterior Hippocampus | Left Cerebellum (Crus 2) | Δ Connectivity x Instructional Methodology | ΔWAIS-IV WMI | -4.784 | 0.291 |
| | | | | ΔWAIS-IV VCI | 2.173 | 0.591 |
| | | | | ΔMean PK Accuracy | 0.093 | **0.026*** |
| | | Right Cerebellum (Crus 1) | | ΔWAIS-IV WMI | -3.826 | 0.344 |
| | | | | ΔWAIS-IV VCI | -0.128 | 0.971 |
| | | | | ΔMean PK Accuracy | -0.022 | 0.569 |
| **Resting State** | | | | | | |
| Main Effect of Time | L. Intermediate Hippocampus | Left Precuneus | Δ Connectivity | ΔWAIS-IV PRI | 1.354 | 0.348 |
| | | | | ΔWAIS-IV WMI | 0.708 | 0.747 |
| | | | | ΔWAIS-IV VCI | 0.463 | 0.812 |
| | | | | ΔWAIS-IV PSI | 1.987 | 0.579 |
| | | Left Precuneus | | ΔWAIS-IV PRI | 0.753 | 0.616 |
| | | | | ΔWAIS-IV WMI | 3.520 | 0.121 |
| | | | | ΔWAIS-IV VCI | 0.157 | 0.938 |
| | | | | ΔWAIS-IV PSI | -3.701 | 0.318 |
| | R. Anterior Hippocampus | Left Lingual Gyrus | | ΔWAIS-IV PRI | 0.344 | 0.838 |
| | | | | ΔWAIS-IV WMI | -1.954 | 0.442 |
| | | | | ΔWAIS-IV VCI | 2.498 | 0.266 |
| | | | | ΔWAIS-IV PSI | -2.246 | 0.589 |
| | | Left Postcentral Gyrus | | ΔWAIS-IV PRI | -2.089 | 0.294 |
| | | | | ΔWAIS-IV WMI | -3.137 | 0.305 |
| | | | | ΔWAIS-IV VCI | 1.722 | 0.520 |
| | | | | ΔWAIS-IV PSI | 4.457 | 0.365 |
| | | Right Lingual Gyrus | | ΔWAIS-IV PRI | 2.543 | 0.179 |
| | | | | ΔWAIS-IV WMI | 3.172 | 0.270 |
| | | | | ΔWAIS-IV VCI | -0.837 | 0.743 |
| | | | | ΔWAIS-IV PSI | 2.149 | 0.648 |

| | | | | | | |
|---|---|---|---|---|---|---|
| Interaction | L. Intermediate Hippocampus | Right Fusiform Gyrus | Δ Connectivity x Instructional Methodology | ΔWAIS-IV PRI | 0.557 | 0.923 |
| | | | | ΔWAIS-IV WMI | 20.735 | **0.020*** |
| | | | | ΔWAIS-IV VCI | 13.417 | 0.086 |
| | | | | ΔWAIS-IV PSI | 15.955 | 0.275 |
| | | Left Superior Frontal Gyrus | | ΔWAIS-IV PRI | -0.966 | 0.908 |
| | | | | ΔWAIS-IV WMI | -11.57 | 0.358 |
| | | | | ΔWAIS-IV VCI | 5.667 | 0.614 |
| | | | | ΔWAIS-IV PSI | -1.997 | 0.923 |

#### Supplemental References

Avants, B., Epstein, C., Grossman, M., & Gee, J. (2008). Symmetric diffeomorphic image registration with cross-correlation: Evaluating automated labeling of elderly and neurodegenerative brain. *Medical Image Analysis*, *12*(1), 26–41. https://doi.org/10.1016/j.media.2007.06.004

Dale, A. M., Fischl, B., & Sereno, M. I. (1999). Cortical Surface-Based Analysis. *NeuroImage*, *9*(2), 179–194.

https://doi.org/10.1006/nimg.1998.0395

Esteban, O., Markiewicz, C. J., Blair, R. W., Moodie, C. A., Isik, A. I., Erramuzpe, A., Kent, J. D., Goncalves, M., DuPre, E., Snyder, M., Oya, H., Ghosh, S. S., Wright, J., Durnez, J., Poldrack, R. A., & Gorgolewski, K. J. (2019). fMRIPrep: A robust preprocessing pipeline for functional MRI. *Nature Methods*, *16*(1), 111–116. https://doi.org/10.1038/s41592-018-0235-4

Greve, D. N., & Fischl, B. (2009). Accurate and robust brain image alignment using boundary-based registration. *NeuroImage*, *48*(1), 63–72. https://doi.org/10.1016/j.neuroimage.2009.06.060

Huntenburg, J. M. (2014). Evaluating Nonlinear Coregistration of BOLD EPI and T1w Images. *Berlin: Freie Universität*.

Jenkinson, M., Bannister, P., Brady, M., & Smith, S. (2002). Improved Optimization for the Robust and Accurate Linear Registration and Motion Correction of Brain Images. *NeuroImage*, *17*(2), 825–841. https://doi.org/10.1006/nimg.2002.1132

Klein, A., Ghosh, S. S., Bao, F. S., Giard, J., Häme, Y., Stavsky, E., Lee, N., Rossa, B., Reuter, M., Chaibub Neto, E., & Keshavan, A. (2017). Mindboggling morphometry of human brains. *PLOS Computational Biology*, *13*(2), e1005350. https://doi.org/10.1371/journal.pcbi.1005350

Reuter, M., Rosas, H. D., & Fischl, B. (2010). Highly accurate inverse consistent registration: A robust approach. *NeuroImage*, *53*(4), 1181–1196. https://doi.org/10.1016/j.neuroimage.2010.07.020

Satopaa, V., Albrecht, J., Irwin, D., & Raghavan, B. (2011). Finding a "Kneedle" in a Haystack: Detecting Knee Points in System Behavior. *2011 31st International Conference on Distributed Computing Systems Workshops*, 166–171. https://doi.org/10.1109/ICDCSW.2011.20

Satterthwaite, T. D., Elliott, M. A., Gerraty, R. T., Ruparel, K., Loughead, J., Calkins, M. E., Eickhoff, S. B., Hakonarson, H., Gur, R. C., Gur, R. E., & Wolf, D. H. (2013). An improved framework for confound regression and filtering for control of motion artifact in the preprocessing of resting-state functional connectivity data. *NeuroImage*, *64*, 240–256. https://doi.org/10.1016/j.neuroimage.2012.08.052

Treiber, J. M., White, N. S., Steed, T. C., Bartsch, H., Holland, D., Farid, N., McDonald, C. R., Carter, B. S., Dale, A. M., & Chen, C. C. (2016). Characterization and Correction of Geometric Distortions in 814 Diffusion Weighted Images. *PLOS ONE*, *11*(3), e0152472. https://doi.org/10.1371/journal.pone.0152472

Tustison, N. J., Avants, B. B., Cook, P. A., Yuanjie Zheng, Egan, A., Yushkevich, P. A., & Gee, J. C. (2010). N4ITK: Improved N3 Bias Correction. *IEEE Transactions on Medical Imaging*, *29*(6), 1310–1320. https://doi.org/10.1109/TMI.2010.2046908

Wang, S., Peterson, D. J., Gatenby, J. C., Li, W., Grabowski, T. J., & Madhyastha, T. M. (2017). Evaluation of Field Map and Nonlinear Registration Methods for Correction of Susceptibility Artifacts in Diffusion MRI. *Frontiers in Neuroinformatics*, *11*. https://doi.org/10.3389/fninf.2017.00017

Yang, H., Zhang, H., Di, X., Wang, S., Meng, C., Tian, L., & Biswal, B. (2021). Reproducible coactivation patterns of functional brain networks reveal the aberrant dynamic state transition in schizophrenia. *NeuroImage*, *237*, 118193. https://doi.org/10.1016/j.neuroimage.2021.118193

Zhang, Y., Brady, M., & Smith, S. (2001). Segmentation of brain MR images through a hidden Markov random field model and the expectation-maximization algorithm. *IEEE Transactions on Medical Imaging*, *20*(1), 45–57. https://doi.org/10.1109/42.906424
